# Mechanical compartmentalization of the intestinal organoid enables crypt folding and collective cell migration

**DOI:** 10.1101/2020.09.20.299552

**Authors:** Carlos Pérez-González, Gerardo Ceada, Francesco Greco, Marija Matejcic, Manuel Gómez-González, Natalia Castro, Sohan Kale, Adrián Álvarez-Varela, Pere Roca-Cusachs, Eduard Batlle, Danijela Matic Vignjevic, Marino Arroyo, Xavier Trepat

**Author notes:** Correspondence to: Xavier Trepat, PhD, ICREA Research Professor, Institute for Bioengineering of Catalonia, Ed. Hèlix, Baldiri i Reixac, 15-21, Marino Arroyo, PhD, Professor, Universitat Politècnica de Catalunya, Danijela Matic Vignjevic, PhD, Research Director INSERM, Institut Curie. These authors contributed equally to this work (Carlos Pérez-González, Gerardo Ceada).

## Abstract

Intestinal organoids capture essential features of the intestinal epithelium such as folding of the crypt, spatial compartmentalization of different cell types, and cellular movements from crypt to villus-like domains. Each of these processes and their coordination in time and space requires patterned physical forces that are currently unknown. Here we map the three-dimensional cell-ECM and cell-cell forces in mouse intestinal organoids grown on soft hydrogels. We show that these organoids exhibit a non-monotonic stress distribution that defines mechanical and functional compartments. The stem cell compartment pushes the ECM and folds through apical constriction, whereas the transit amplifying zone pulls the ECM and elongates through basal constriction. Tension measurements establish that the transit amplifying zone isolates mechanically the stem cell compartment and the villus-like domain. A 3D vertex model shows that the shape and force distribution of the crypt can be largely explained by cell surface tensions following the measured apical and basal actomyosin density. Finally, we show that cells are pulled out of the crypt along a gradient of increasing tension, rather than pushed by a compressive stress downstream of mitotic pressure as previously assumed. Our study unveils how patterned forces enable folding and collective migration in the intestinal crypt.

---

Most epithelial tissues that line internal and external surfaces of the animal body perform their diverse physiological functions in a continuous state of self-renewal^1^. In the intestinal epithelium, rapid self-renewal is enabled by stem cells that reside at the bottom of highly curved invaginations called crypts, where they coexist with secretory Paneth cells^2^. To maintain homeostasis, stem cells constantly divide, giving rise to new cells that proliferate further at the transit amplifying zone, differentiate, and migrate to the tip of finger-like protrusions called villi, where they are extruded into the intestinal lumen^3,4^. These processes are recapitulated by intestinal organoid models, which contain the main cell types of the intestinal epithelium, compartmentalize these cell types into functional units, fold into a crypt-like geometry, and capture essential features of cell proliferation, differentiation, motility and extrusion^5–8^.

The intestinal epithelium self-renews while remaining folded into crypts and villi. Folding of the intestinal surface into villi has been studied in different systems and is attributed to compressive stresses generated by the mesenchyme or the smooth muscle^9–11^. By contrast, how the crypt folds during morphogenesis and how folding is maintained during homeostasis is not well understood. Candidate folding mechanisms include buckling as a consequence of either increased mitotic pressure^12–15^, differentials in actomyosin forces between epithelial compartments^16^, or planar cellular flows^17^. Additionally, crypt folding could arise from bending by apical constriction^18,19^, basal expansion^20^ or transepithelial differences in osmotic pressure^21,22^. Importantly, the cellular forces that fold the epithelium must also enable other mechanical functions such as collective movement from crypt to villus^4^. How forces are distributed in the intestinal epithelium to drive its diverse mechanical functions is unknown. Using the mouse intestinal organoid as a model system, here we provide high-resolution dynamic maps of cell-ECM and cell-cell forces exerted by the intestinal epithelium. Our experiments and computational model reveal that the intestinal epithelium is organized in mechanical compartments in which patterned forces drive folding by apical constriction while enabling cell migration along tensile gradients.

To study the mechanics of intestinal crypts, we established a bottom up approach (Fig. 1a). Rather than beginning with fully folded organoids surrounded by a 3D ECM, we allowed isolated mouse intestinal crypts to spread on soft and flat substrates, and then studied whether and how the resulting 2D sheets refolded and retained homeostatic functions such as proliferation, differentiation, migration and extrusion. Mouse intestinal crypts readily spread on soft (5 kPa) polyacrylamide gels coated with collagen-I and laminin-1, in a process akin active tissue wetting (Supplementary Video 1)^23^. A few days after crypt spreading, the substrate was covered by a continuous monolayer that showed a clear compartmentalization of the main cell types of the intestinal epithelium (Fig. 1b, Extended Data Fig.1). Stem cells (labelled with Lgr5 and Olfm4) and Paneth cells (labelled with lysozyme) were intermingled in a roughly circular compartment devoid of any other cell type. This stem cell compartment was surrounded by a ring of highly proliferative cells (positive for Ki67) elongated in the direction parallel to the crypt contour. At the outer edge of this ring, cells expressed differentiation marker cytokeratin-20 and increased their spreading area. These experiments indicate that the intestinal epithelium can be cultured on soft 2D substrates for several days while maintaining the spatial segregation of its different cell types.

**Fig. 1.**
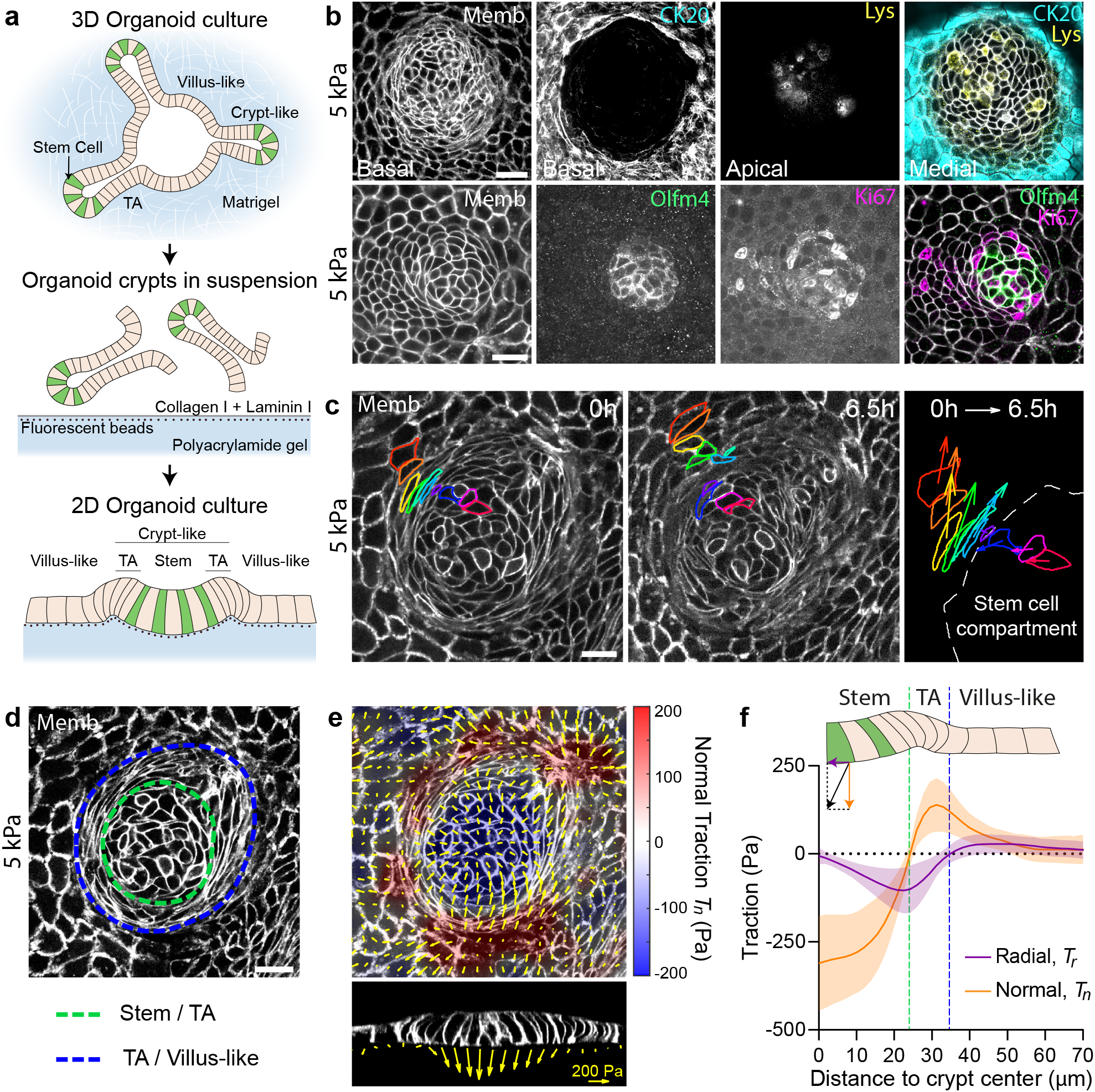
Tractions exerted by intestinal organoids define mechanical compartments. **a**, Preparation of mouse intestinal organoids on 2D soft substrates. **b**, Organoids expressing membrane targeted tdTomato (Memb, basal plane) stained for cytokeratin 20 (CK20, basal plane), Lysozyme (Lys, apical plane), Olfm4 (basal plane) and Ki67 (basal plane). Scale bar, 20 µm. **c**, Displacements of representative cells over 6.5h. Each color labels one cell. Note that one cell divided (green). Right: displacement vector of each cell. Scale bar, 20 µm. See also Supplementary Video 3. **d**, Illustration of the boundaries between the stem cell compartment and the transit amplifying zone (green) and between the transit amplifying zone and the villus-like domain (blue). Scale bar, 20 µm. **e**, Top: 3D traction maps overlaid on a top view of an organoid. Yellow vectors represent components tangential to the substrate and the color map represents the component normal to the substrate. Bottom: lateral view along the crypt horizontal midline. Yellow vectors represent tractions. Scale vector, 200 Pa. **f**, Circumferentially averaged normal tractions T_*n*_ (orange) and radial tractions T_*r*_ (purple) as a function of the distance to the crypt center. Blue and green dashed lines indicate the radii where *T*_*n*_ and *T*_*r*_ are zero, which closely correspond to the boundaries between functional compartments illustrated in **d**. Data are represented as mean ± SD of N=36 crypts from 7 independent experiments.

We next asked whether cells in each compartment retained their *in vivo* homeostatic functions. Time-lapse imaging showed that stem cells divided frequently and that some daughter cells entered the highly proliferative zone, where they divided further (Fig. 1b, Supplementary Videos 2, 3). Upon leaving this zone, cells continued to move radially out well into the differentiated zone, where they ultimately died and were extruded (Supplementary Video 4). Given the analogy between organoid monolayers *in vitro* and the intestinal epithelium *in vivo*, we will adopt the following nomenclature: the region containing only stem cells and Paneth cells will be called the stem cell compartment; the region containing highly proliferative elongated cells will be called the transit amplifying zone (TA); and the region containing differentiated cells will be called the villus-like domain (Fig. 1b,d).

We used traction microscopy to map the 3D traction forces exerted by the epithelial monolayer on its underlying substrate^22,24^. Traction maps and their circumferential averages revealed systematic mechanical patterns that were radially symmetric about the crypt center (Fig. 1d, e). We decomposed these traction patterns into a normal component (*T*_*n*_, perpendicular to the substrate) and a radial component (*T*_*r*_, parallel to the substrate). At the stem cell compartment, normal tractions were negative, indicating that stem cells and Paneth cells pushed the substrate downward (Fig. 1e, f). The magnitude of normal tractions was highest at the crypt center, decreased radially away from it, and vanished at the margin of the stem cell compartment. At the transit amplifying zone, normal tractions became positive, indicating that cells in this region pulled the substrate upward. After peaking at the transit amplifying zone, normal tractions progressively decreased and vanished in the villus-like domain except in a few areas where the monolayer had delaminated to form domes (Extended Data Fig 2, Supplementary Video 4). In these domes, which can be loosely interpreted as villus-like protrusions, tractions pointed towards the substrate, showing that luminal fluid was pressurized^22^. The radial component of the traction fields also followed a non-monotonic behavior. This component was negligible at the center of the stem cell compartment and became progressively negative (i.e., pointing inward) towards its border, where it peaked near the transit amplifying zone. At the outer border of this zone, radial tractions vanished and then became slightly but systematically positive (i.e., pointing outward) in the villus-like domain. Overall, these measurements reveal that the intestinal epithelium generates non-monotonic traction fields on the substrate, defining distinct mechanical compartments that colocalize with functional compartments; the stem cell compartment pushes downward, the transit amplifying zone pulls upward and shears inward, and the villus-like domain shears outward.

**Fig. 2.**
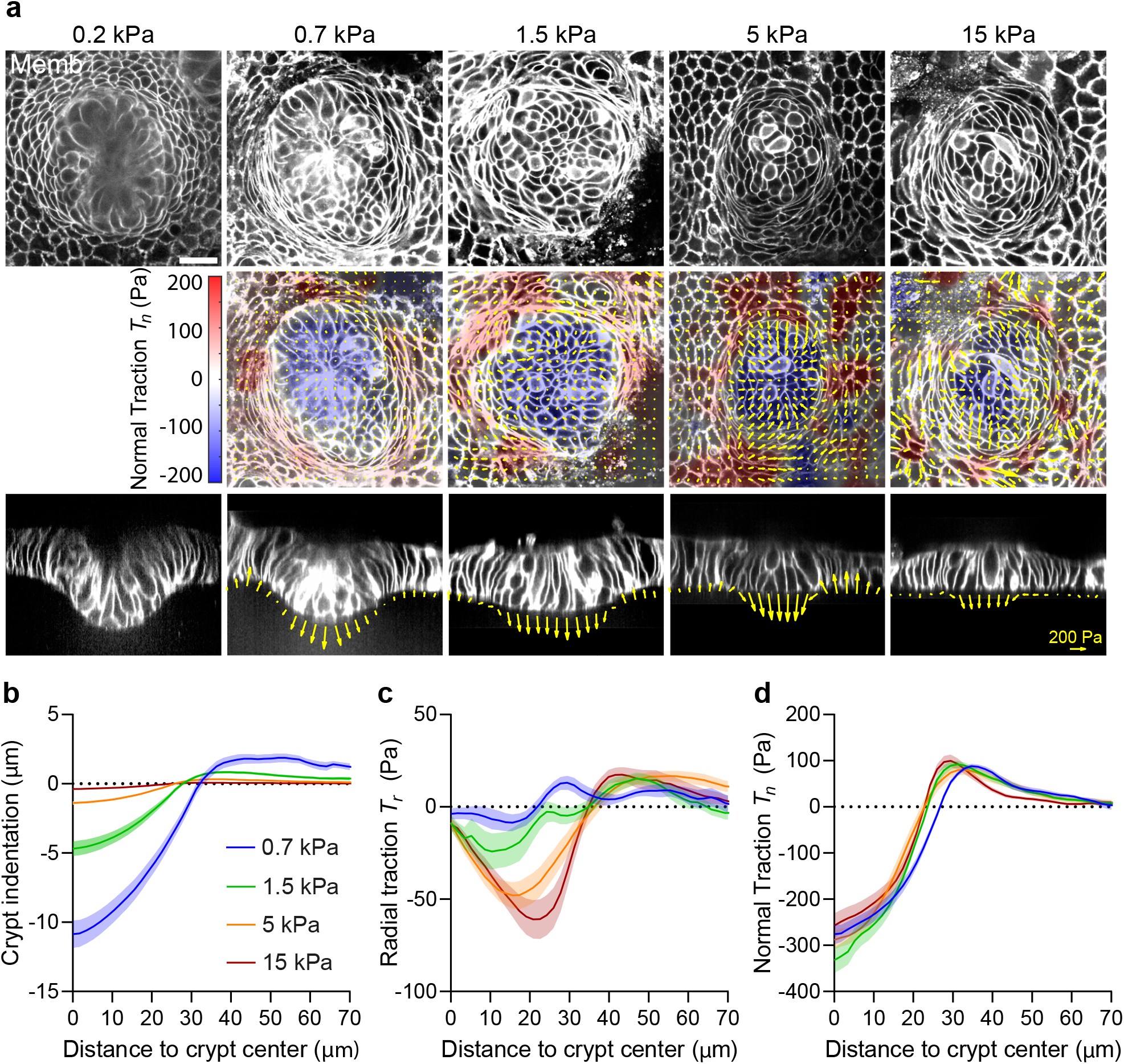
A stiffness-independent normal traction folds the crypt. **a**, Top: single confocal plane of representative crypts on substrates of increasing stiffness. Center: 3D traction maps. Yellow vectors represent components tangential to the substrate and the color map represents the component normal to the substrate. Bottom: lateral view along the crypt midline. Yellow vectors represent tractions. Scale bar, 20 µm. Scale vector, 200Pa. **b-d**, Crypt indentation (**b**), radial traction (**c**) and normal traction (**d**) as a function of the distance to the crypt center for substrates of different stiffness. Data are represented as mean ± SEM of N=14 (0.7 kPa), 12 (1.5 kPa), 36 (5 kPa) and 30 (15 kPa) crypts from, 3, 3, 7 and 4 independent experiments, respectively.

We next asked how these mechanical compartments control the shape and function of the crypt and its transition towards the villus-like domain. We begin investigating the mechanism that drives crypt folding. Our measurements of normal tractions (Fig. 1e, f) suggest that the crypt folds by pushing the stem cell compartment towards the substrate, but folding was frustrated because the underlying substrate was too stiff. Thus, we investigated whether a sufficiently soft substrate would enable the stem cell compartment to adopt the folded shape characteristic of 3D cultured organoids and *in vivo* crypts. To study the interplay between substrate stiffness, tractions and folding, we cultured intestinal organoids on substrates of varying stiffness, spanning nearly two orders of magnitude (0.2, 0.7, 1.5, 5, 15kPa, Young’s Modulus). On the stiffest substrate (15kPa), the stem cell compartment bulged out of the monolayer (Fig. 2a). By contrast, when organoids were grown on progressively soft substrates, they showed an increasingly pronounced folding (Fig. 2a, b). We measured cell tractions for each substrate stiffness except for the 0.2kPa substrates, which displayed extreme deformations and hydrogel creasing instabilities that prevented a sufficiently accurate calculation (Fig. 2a). The radial traction component exhibited pronounced changes with substrate stiffness (Fig. 2c). As monolayers progressively folded, this component nearly vanished at the stem cell compartment, while remaining weak but positive in the villus-like domain. By contrast, the normal traction component was independent of substrate stiffness (Fig. 2d). Thus, the traction component parallel to the substrate was mechanosensitive, as commonly observed in most cell types^25–27^, but the normal component was not, suggesting that the different force components are generated by different cellular structures.

The observed traction patterns and monolayer geometry can be explained by two classes of folding mechanisms. The first class relies on differentials in mitotic pressure^12–14^, which can induce a buckling instability that pushes the stem cell compartment towards the substrate. A second class is based on differentials in myosin contractility, either across the monolayer plane^16^ or along the apicobasal axis^18–20^. To discriminate between these two classes of mechanisms we treated cells with blebbistatin. Upon addition of this specific inhibitor of myosin II ATPase activity, all traction forces vanished (Fig. 3a-c) and the monolayer flattened (Fig. 3d, e, Supplementary Video 5). The elongated basal shape of cells at the transit amplifying zone, which was unaffected by the stiffness of the gel, was largely lost by this treatment. When blebbistatin was washed out, traction patterns reemerged, the elongated morphology was recovered, and the monolayer refolded (Fig. 3a-c). Thus, tuning myosin activity is sufficient to reversibly control the shape and mechanics of the intestinal epithelial monolayer.

**Fig. 3.**
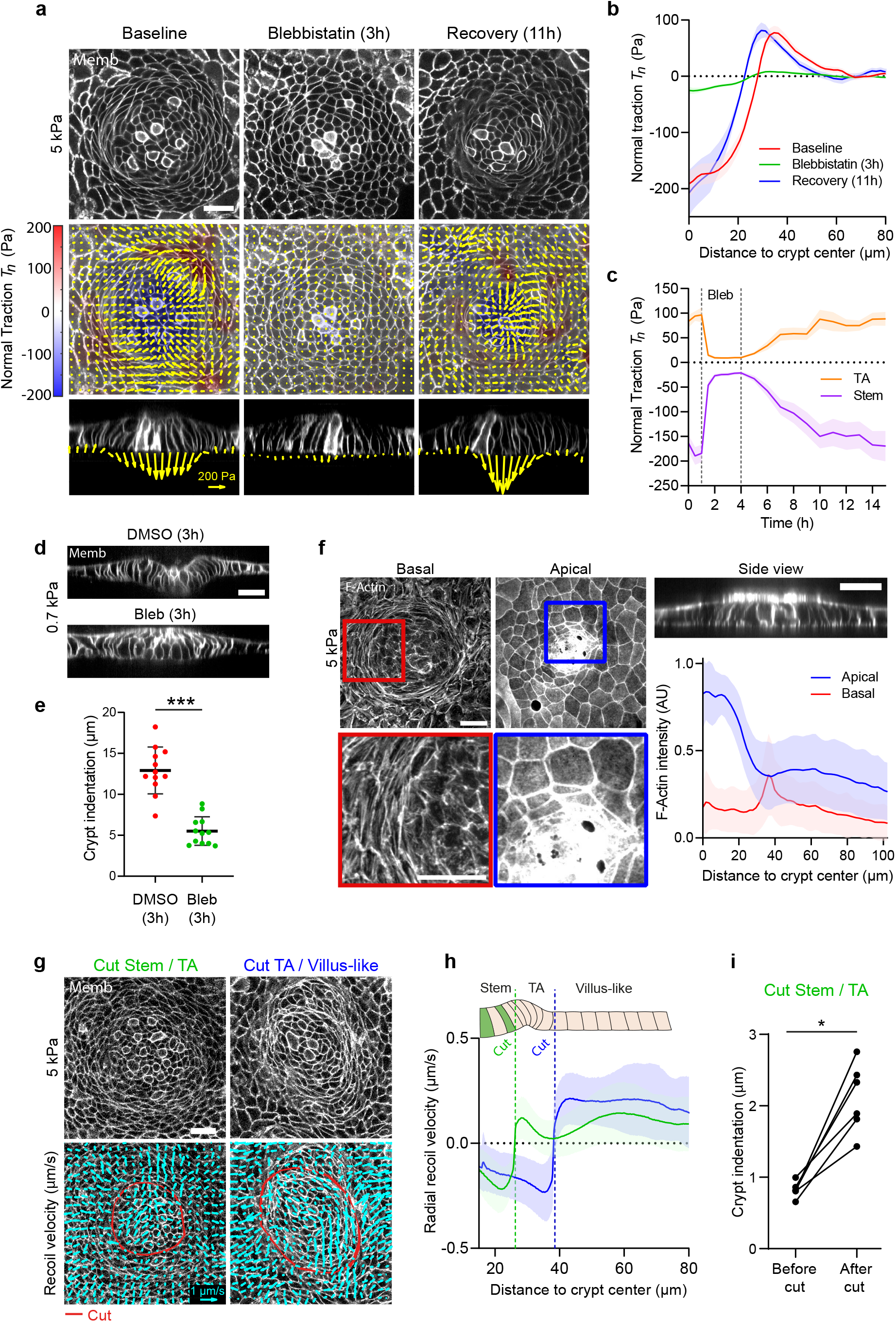
The crypt folds under tension generated by myosin II. **a**, Traction maps under baseline conditions (left), after 3h of treatment with blebbistatin (center), and after 11h of blebbistatin washout (right). Top: membrane targeted tdTomato (medial confocal plane). Center: 3D traction maps. Yellow vectors represent components tangential to the substrate and the color map represents the component normal to the substrate. Bottom: lateral view of the organoids along the crypt midline. Yellow vectors represent tractions. Scale bar, 20 µm. Scale vector, 200Pa. **b**, Normal traction as a function of the distance to crypt center before, during, and after blebbistatin treatment. Data are represented as mean ± SEM of N=11 samples from 3 independent experiments. **c**, Time evolution of normal traction for the stem cell compartment and the transit amplifying zone before, during and after blebbistatin treatment. Data are represented as mean ± SEM of N=11 samples from 3 independent experiments. **d**, Lateral views of crypts seeded on 0.7kPa gels and treated with DMSO (top) or blebbistatin (bottom) for 3h. Scale bar, 20 µm. **e**, Indentation at the center of the crypt in cells treated with DMSO or blebbistatin for 3h. Mean ± SD of N=12 crypts from 2 independent experiments. *** indicates p<0.001 (two-tailed t-test). **f**, F-actin (phalloidin) staining of crypts seeded on 5kPa gels. Left and center: projections of the basal (left) and apical (center) F-actin intensity of a representative crypt. Top right: Lateral view of the same crypt. Bottom right: Radial distribution of the apical (blue) and basal (red) F-actin intensity as a function of the distance to the crypt center. Scale bars, 20 µm. Data are presented as mean ± SD of N=37 crypts from 4 independent experiments. **g**, Bottom: Recoil velocity maps immediately after ablation of two crypts along the red lines. Left: a cut inside the TA. Right: a cut outside the TA. Scale vector, 1 µm/s. Top: the two crypts before ablation. **h**, Radial recoil velocity as a function of distance to crypt center for cuts between stem cell compartment and transit amplifying zone (green) and between transit amplifying zone and villus-like domain (blue). Data are represented as mean ± SD of N=14 (cut Stem / TA) and 11 (cut TA / Villus-like) crypts from 5 independent experiments. **i**, Indentation at the center of the crypt before and after cutting at the interface between the stem cell compartment and the TA. N=6 crypts from 2 independent experiments. * indicates p<0.05 (Wilcoxon paired test).

To study how the actomyosin cytoskeleton drives crypt folding, we measured the localization of actin and myosin across the monolayer. F-actin stainings (phalloidin) and live imaging of organoids expressing myosin IIA-eGFP revealed an actomyosin accumulation at the apical surface of the stem cell compartment (Fig. 3f, Extended Data Fig. 3). In addition, they showed that the actomyosin cytoskeleton formed a basal ring of circumferential fibers under the elongated cells at the transit amplifying zone. This actomyosin distribution mirrors the one observed in the developing crypt in vivo^18^ and suggests two potential mechanisms that are not mutually exclusive. The first one is that myosin differentials across the apicobasal cell axis drive monolayer bending through apical constriction or basal relaxation. The second one is that supracellular contraction of the transit amplifying zone compresses radially the stem cell compartment to induce its buckling. To assess the contribution of bending *vs* buckling, we reasoned that whereas both mechanisms could result in similar cell-substrate traction patterns, they involve cell-cell stresses of opposite sign. In the bending scenario, cells in the stem cell compartment should pull on each other differentially along the apicobasal axis to indent the substrate, thereby generating an apical tensile stress. By contrast, in the buckling scenario, cells in the stem cell compartment should push on each other as a consequence of the compressive stress generated by the contractile ring at the transit amplifying zone.

To measure the sign of the stress field, we performed circular laser ablations along the internal or external boundaries of the transit amplifying zone. In both cases, ablations induced a radial recoil on both sides of the cuts (Fig. 3g, Supplementary Videos 6, 7). Recoils were asymmetric and showed non-monotonic recoil velocity fields, indicating that monolayer friction and viscosity follow a complex spatial distribution that prevents a straightforward readout of relative tensions from recoil dynamics (Fig. 3h). However, the sole fact that monolayers recoiled on both sides of the cuts implies that the crypt is under tension.

Moreover, after cutting the monolayer at the inner boundary of the transit amplifying zone, the stem cell compartment remained folded and it even increased its indentation of the substrate (Fig. 3i). This set of experiments rules out the possibility that the crypt forms through compressive buckling driven by myosin differentials between crypt compartments. It also rules out other compressive buckling scenarios such as differentials in mitotic pressure. Instead, it shows that the stem cell compartment bends by apical constriction and that the supracellular actomyosin ring located basally at the transit amplifying zone is not required for folding, further highlighting the mechanical compartmentalization of the crypt.

We next investigated how cell shape evolves between compartments. To this aim, we segmented individual cell shape in crypts grown on stiff (15kPa) and soft (0.7kPa) substrates. Cell shapes were highly heterogeneous, but averages over all cells and crypts revealed consistent morphometric patterns (Fig. 4a, b). Both on stiff and soft substrates, the apical area of stem cells was smaller than the basal one (Fig. 4c, d). Paneth cells showed the opposite behavior (Extended Data Fig. 4), indicating that stem but not Paneth cells are responsible for apical constriction of the stem cell compartment. Both cell types were taller than differentiated cells (Fig. 4e) and displayed an apicobasal tilt towards the center of the crypt that peaked at the boundary between the stem cell compartment and the transit amplifying zone (Fig. 4f). Upon reaching this boundary, the differences between apical and basal area disappeared and cells became basally elongated along the circumferential direction (Fig. 4g), parallel to the supracellular actomyosin ring. Treatment with blebbistatin for 3h reduced significantly the tilt at the stem cell compartment and the basal aspect ratio of every cell type at the crypt (Extended Data Fig. 5).

**Fig. 4.**
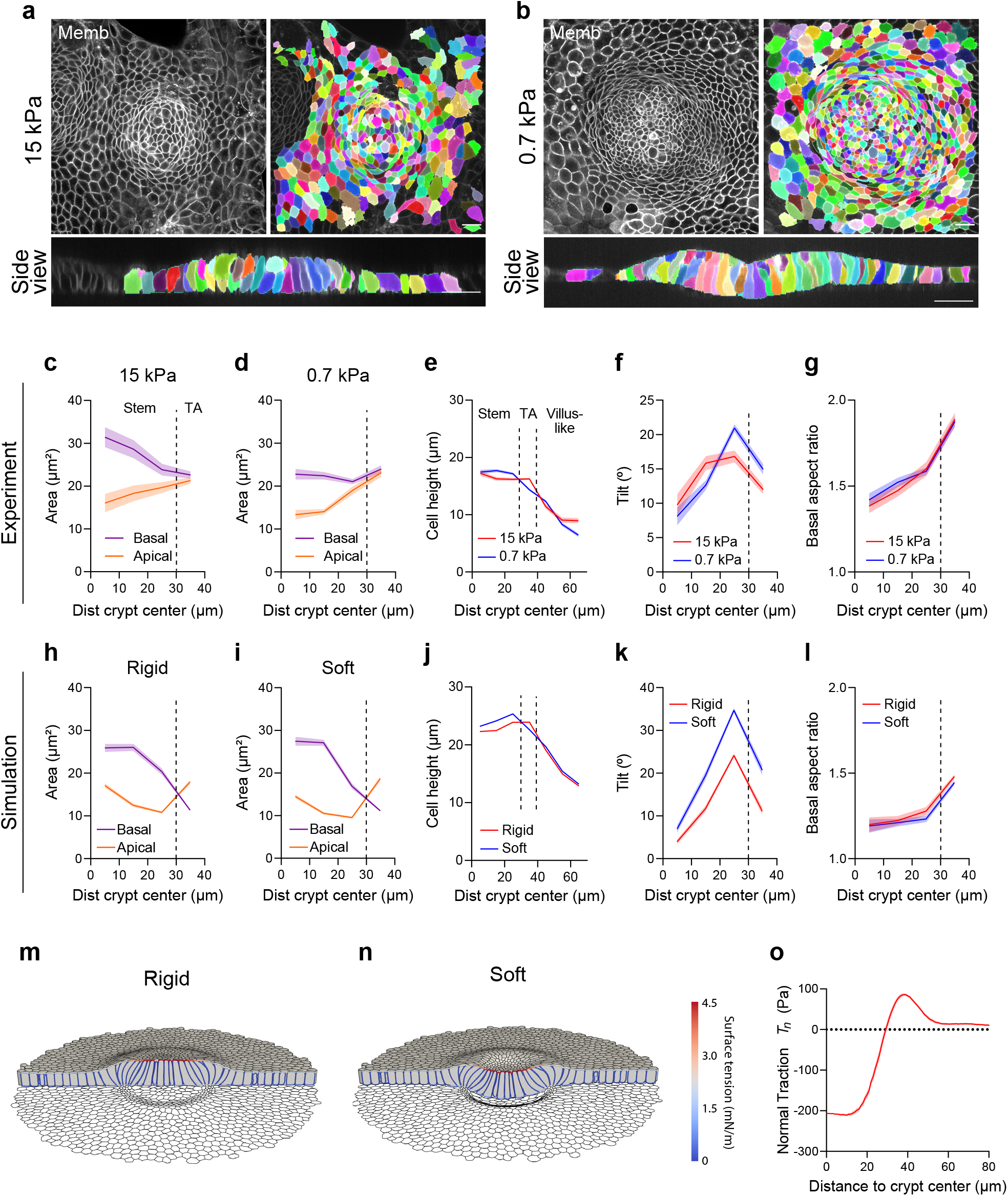
Crypts fold through apical constriction. **a-b**, 3D Crypt segmentation on stiff (**a**, 15kPa) and soft (**b**, 0.7kPa) substrates. **c-d**, Apical and basal area as a function of the distance to the crypt center on stiff (**c**, 15 kPa) and soft (**d**, 0.7 kPa) substrates. Paneth cells were excluded for the analysis (see Extended Data Fig. 4). Data are represented as mean ± SEM. From center to edge bins, N=77, 198, 339 and 307 cells for soft substrates and 32, 61, 102 and 230 cells for stiff substrates. N=3 independent crypts per stiffness from 2 (0,7kPa) and 3 (15kPa) independent experiments. **e-g** Cell height (**e**), apicobasal tilt (**f**) and basal aspect ratio (**g**) as a function of distance to crypt center on soft and stiff substrates. Paneth cells were excluded from the analysis of the tilt and aspect ratio (see Extended Data Fig. 4). Vertical dashed line indicates boundary between stem cell compartment, the transit amplifying zone and the villus-like domain (in **e**). Crypts are the same as in panel **c**-**d**. For cell height profiles, from center to edge bins, N= 77, 198, 339, 307, 242, 242 and 242 cells for soft substrates and 32, 61, 102, 230, 242, 242 and 242 cells for stiff substrates. **h-i**, Simulated apical and basal areas as a function of distance to crypt center on a stiff (**h**) and soft (**i**) substrate. Data are presented as mean ± SEM. From center to edge bins, N= 13, 35, 49 and 216 simulated cells for soft substrate and N= 13, 31, 52 and 217 simulated cells for rigid substrate. **j-l**, Simulated cell height (**j**), apicobasal tilt (**k**) and basal aspect ratio (**l**) as a function of distance to crypt center on soft and stiff substrates. Crypts are the same as in **h-i. m-n**, 3D vertex model of an intestinal monolayer adhered to a stiff (**i**) and soft (**j**) substrate. The colors in the cell outlines indicate the magnitude of surface tension, prescribed according to measured F-actin intensity. The lateral dimension of the monolayer is 150 µm and the cell height in the villus-like region is 10 µm. **o**, Simulated normal traction in the 3D vertex model of the intestinal monolayer. To be compared with the experimental values in Fig. 1f and Fig. 2d.

**Fig. 5.**
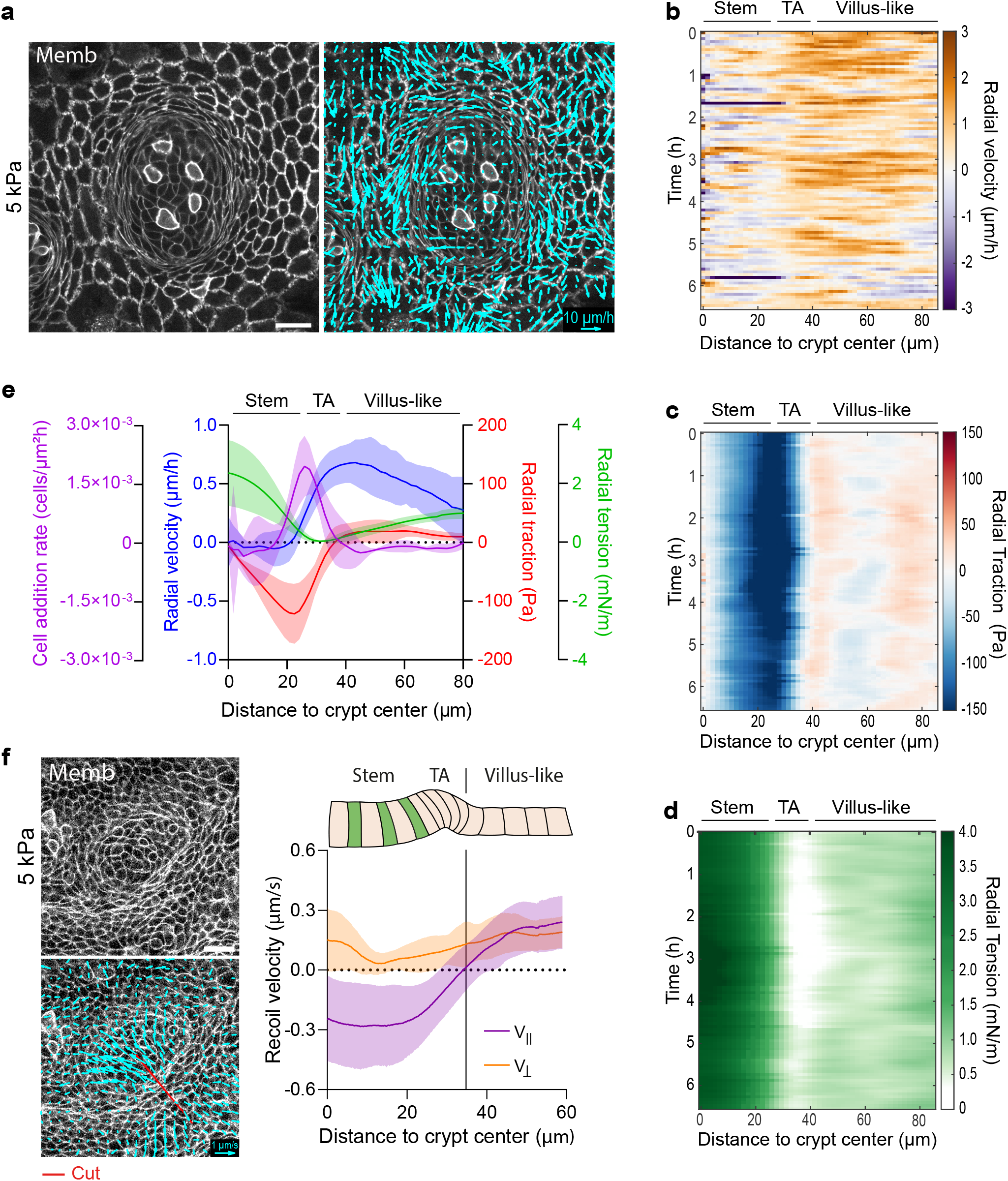
Cells are dragged out of the crypt towards the villus-like domain. **a**, Representative velocity maps (right) overlaid on membrane-targeted tdTomato signal (shown in left panel), Scale bar, 20 µm. Scale vector, 10 µm/h. **b-d**, Representative kymographs showing the circumferentially averaged radial velocity (**b**), radial traction (**c**) and radial tension (**d**) during 6.5hours. **e**, Average radial profiles of traction, tension, velocity and cell addition rate. Data are represented as mean ± SD of N=5 crypts from 3 independent experiments. Note that for an unbounded monolayer, MSM computes stress up to a constant (Supplementary Note 2). Since laser cuts indicated tension everywhere in the monolayer, this constant was arbitrarily setup so that the minimum tension was zero. **f**, Bottom left: Recoil velocity maps immediately after laser ablation along the crypt-villus axis. Top left: the crypt before ablation. Right: recoil velocity parallel and perpendicular to the cut as a function of distance to the crypt center. Scale vector, 1 µm/s. Scale bar, 20 µm. Data are represented as mean ± SD of N=11 crypts from 3 independent experiments.

To investigate the link between measured tissue shape, cell shape, cell-ECM tractions, and actomyosin localization we developed a 3D vertex model^28–30^ of the crypt allowing for complex cell shapes and coupled to a soft hyperelastic substrate (Supplementary Note 1, Extended Data Figs. 6, 7). We considered a uniform flat monolayer with a pattern of cell surface tensions as suggested by the measured distribution of cortical components (Fig. 3f, Extended Data Fig. 3). Specifically, we prescribed apical and basal surface tensions with a profile matching the measured F-actin density (Fig. 3f). We prescribed much smaller lateral surface tensions as suggested by the observed smoothness of apical and basal surfaces near lateral cell junctions (Supplementary Note 1). We then allowed the monolayer to equilibrate its shape while adhering to a rigid substrate. We repeated the computational analysis on a soft substrate under the exact same surface tension pattern. The model was able to recapitulate all morphological features of the monolayer, both at the cell and tissue scales, including cell height, shape and monolayer folding for both soft and stiff substrates (Fig. 4m, n). The model captured apical constriction (Fig. 4h, i) and increased cell height (Fig. 4j) at the stem cell compartment, apicobasal cell tilt (Fig. 4k), and basal tangential elongation at the transit amplifying zone (Fig. 4l). Besides cell and tissue morphology, the model also predicted the distribution of normal cell-substrate tractions (Fig. 4o). The magnitude of the measured normal tractions allowed us to estimate the magnitude of the required apical surface tensions, which we found in the order of 4.6 mN/m. This remarkable agreement shows that a strongly stereotyped contractility pattern in the crypt, with 10-fold differences in surface tension, can explain its stiffness-dependent shape and normal traction patterns. We note, however, that this mechanical picture is static, and does not explain dynamic features of the epithelial monolayer such as cell movements from crypt to villus-like domains, which we set out to study next.

Cell movements at the villus were recently shown to be driven by active migration^31^, but mechanisms driving movements from crypt to villus remain unknown. In fact, the widely assumed mechanism for such movements, a pushing force arising from compression downstream of mitotic pressure^32,33^, is incompatible with our observation that the transit amplifying zone is under tension. To study the mechanisms underlying cell movement, we mapped cell velocity in the organoid monolayers on 5kPa gels using particle imaging velocimetry (PIV). Velocity maps revealed strong spatial fluctuations, characterized by clusters of fast-moving cells surrounded by nearly immobile ones (Fig. 5a, Supplementary Video 8). To average out these fluctuations and unveil systematic spatiotemporal patterns, we computed the average radial velocity as a function of the distance from crypt center, and represented this average as a kymograph (Fig. 5b). Kymographs revealed that cell velocity was intermittent in time, with periods of slow movement alternating with rapid ones. Upon averaging the kymographs over time and experimental repeats, we found a clear velocity pattern. Radial velocity was negligible in the central region of the stem cell compartment, begun to increase near the transit amplifying zone and peaked at the boundary with the villus-like domain (Fig. 5b, e). By also measuring the cell density profile (Extended Data Fig. 8), the radial velocity profile allowed us to establish an equation for balance of cell number and thus quantify the cell addition rate (Supplementary Note 2). This quantification confirmed that most divisions occurred at the outer boundary of the stem cell compartment and at the transit amplifying zone (Fig. 5e).

To understand how this spatial distribution of cell velocity arises from cellular forces, we turned to kymographs of radial cell-ECM tractions and radial cell-cell monolayer tension measured with Monolayer Stress Microscopy (MSM, Supplementary Note 3)^34^. Traction and tension kymographs were stationary, confirming that our 2D organoids are in steady state within our window of observation (Fig. 5c, d). When kymographs were averaged over time and experimental repeats, radial traction displayed the three compartments already shown in Fig. 1e; in the stem cell compartment radial tractions pointed inwards, in the transit amplifying zone they vanished, and in the villus-like domain they pointed outwards (Fig. 5e). By contrast, kymographs of cell-cell tension showed a minimum at the transit amplifying zone flanked by two tension gradients that build-up towards the center of the crypt and towards the villus-like domain (Fig. 5d, e). To further substantiate this tensional compartmentalization, we resorted again to laser ablation, but this time we cut the monolayer radially. Unlike commonly observed in laser ablation experiments, recoil following ablation was predominantly parallel to the cut, rather than perpendicular to it (Fig. 5f, Supplementary Video 9). Radial recoil was minimum at the transit amplifying zone and increased on either side of it, supporting that this zone separates two compartments of opposing tensile gradients in agreement with our data obtained with MSM.

Together, our results lead to the following mechanical picture. The stem cell compartment and the villus-like domain define two largely independent mechanical compartments. At the center of the stem cell compartment, net cell movements are negligible and apical constriction folds the crypt while generating a negative tensile gradient. At the transit amplifying zone, cells accelerate and elongate basally in the circumferential direction. In this zone, radial tension reaches a minimum value that coincides with the region of highest division rate, suggesting that proliferation contributes to lower tension at the transit amplifying zone. As this zone transitions towards the villus-like differentiated area, a second tension gradient builds up. Cells migrate up this gradient in a migration mode in which cell velocity is parallel to cell-substrate traction, rather than antiparallel to it as has been previously observed at the leading edge of advancing monolayer sheets^35,36^. This result establishes that cells leave the transit amplifying zone dragged by cells located further into the villus-like domain, much as the trailing edge of a cell cluster is dragged by the leading edge^37^. Thus, cells exit the crypt through an unanticipated mode of migration that can be understood as a multicellular trailing edge that is constantly being replenished with cells through proliferation at the transit amplifying zone. The mechanical picture portrayed here is nontrivial and challenges previous views on how mechanics influences epithelial tissue patterning. Indeed, previous studies in differentiated epithelial cell lines found that tension triggers proliferation^38–40^, whereas here we found that proliferation is highest where tension is lowest. Previous work also reported that compression triggers extrusion^41,42^, whereas here we found frequent extrusion in areas of high tension. Our work provides a new experimental basis to reformulate current models of how cellular forces influence epithelial self-renewal and homeostasis^31,43^.

The field of organoid biology has revolutionized life sciences by providing accessible systems to study the interplay between tissue fate, form and function. Mechanics plays a central role in this interplay^44–47^, but high-resolution time-evolving maps of the forces that cells within organoids exert on their surroundings have evaded experimental observation. Here we presented mechanically-accessible intestinal organoids that display homeostatic functions, compartmentalization of cell types, and folding into crypt-like geometry. Besides enabling straightforward imaging and versatile control of the mechanical environment, our organoids display an open apical surface^48–51^ and collective migration from crypt to villus-like domains, two features that are not well captured by 3D intestinal organoids surrounded by ECM. Direct access to cell-ECM and cell-cell forces allowed us to identify the mechanisms by which forces exerted in each compartment enable both tissue folding and collective cell migration. Other key processes in intestinal physiology and disease, such as proliferation, differentiation, inflammation, regeneration and malignancy are also likely to be regulated by physical forces. The mechanobiology of these processes is now amenable to quantitative experimental study.

## Methods

### Intestinal crypt isolation and organoid culture

Animal experimentation was approved by the Animal care and Use Committee of Barcelona Science Park (CEEA-PCB) and Animal Welfare Body, Research Centre, Institut Curie. All procedures were carried out in compliance with the European Regulation for the Protection of Vertebrate Animals used for Experimental and other Scientific Purposes (Directive 2010/63). Intestinal crypts from mT/mG^52^, LifeAct-eGFP^53^, MyosinIIA-eGFP^54^, and Lgr5-eGFP-IRES-CreERT2^2^ mice were isolated as described elsewhere^5^. The duodenum was isolated, washed in PBS and cut longitudinally. Villi were mechanically removed with a scalpel. The tissue was then dissociated in HBSS (Gibco) containing 8mM of EDTA (Sigma-Aldrich) for 20 minutes at 4^°^C. After vigorous shaking, the solution was filtered through a 70 µm pore cell strainer (Corning), obtaining the crypt fraction. The dissociation, shaking and filtration steps were repeated three times to increase the isolation’s yield. The final solution was centrifuged at 100xg for 5 minutes at room temperature (RT) and the pellet containing crypts was resuspended in 1:1 Matrigel (Corning): ENR* and seeded in drops of 50 µL in 24 well plates. After 25 min incubation at 37^°^C 5%CO_2_ in an humified atmosphere, the drops were covered with ENR medium*. The organoids were split every 3-4 days. For splitting, organoid-containing drops were mechanically disaggregated by pipetting in PBS supplemented with calcium and magnesium (Sigma-Aldrich) and centrifuged at 100xg at RT for 3.5 minutes. The pellet was resuspended in 1:1 Matrigel: ENR and seeded in drops as explained above.

*ENR medium: DMEM/F-12 (Gibco) supplemented with 2% Antibiotic/Antimycotic (Gibco), 2.5% Glutamax (Gibco), 20ng/mL mouseEGF (Peprotech), 100ng/mL Noggin (Peprotech), 500ng/mL R-spondin1 (R&D Systems), 10ng/mL mouseFGF (Peprotech), 1X B-27 (Gibco) and 1X N-2 (Gibco).

### Polyacrylamide (PAA) gel polymerization

Glass bottom dishes (MatTek) were incubated 10 minutes at RT with Bind-Silane (Sigma-Aldrich) dissolved in ethanol absolute (PanReac) and acetic acid (Sigma-Aldrich) at volume proportions 1:12:1, respectively. After two washes with ethanol absolute, 22.5µL of the polyacrylamide mix (see Supplementary Table 1 for the different recipes used) were added on top of the glass and covered with an 18 mm coverslip. After 1h polymerization at RT, PBS was added, and the coverslips were removed with a scalpel.

**Supplementary Table 1.** Polyacrylamide mix

| Stiffness (kPa) | Acrylamide (BioRad) % | Bis-acrylamide (BioRad) % | Beads** % solids | Ammonium persulphate (Sigma-Aldrich) % | Tetramethylethylenediamine (Sigma-Aldrich) % |
| --- | --- | --- | --- | --- | --- |
| 0.2 | 3 | 0.03 | 0.04 | 0.5 | 0.05 |
| 0.7 | 4 | 0.03 | 0.04 | 0.5 | 0.05 |
| 1.5 | 5 | 0.04 | 0.04 | 0.5 | 0.2 |
| 5 | 7.46 | 0.044 | 0.04 | 0.5 | 0.05 |
| 15 | 7.5 | 0.16 | 0.04 | 0.5 | 0.05 |

\*\* For traction measurements we used FluoSpheres of 0.2µm diameter (Thermofisher). For immunostainings, we used CML Latex Beads of 0.2µm diameter (Thermofisher).

### PAA gel functionalization and ECM micropatterning

Organoid monolayers were confined in large circular ECM micropatterns (>900mm). For this, PAA gels were functionalized with 2mg/mL of Sulpho-SANPAH (Cultek) irradiated for 7.5 minutes with UV light (365nm). Gels were then washed twice with HEPES 10mM (Gibco). For ECM micropatterning, Polydimethylsiloxane (PDMS) stencils with circular openings were used^23^. PDMS stencils were incubated with Pluronic acid F127 2% in PBS (Sigma-Aldrich) for 1 hour. Then, they were washed twice with PBS and allowed to dry at RT for 20 minutes. Stencils were carefully placed on top of the functionalized PAA gels. A solution of 250 µg/mL rat tail type I Collagen (First Link UK) and 100 µg/mL Laminin (SigmaAldrich) dissolved in PBS was added on top of the PDMS stencils and incubated overnight at 4^°^C. Finally, the ECM solution was aspirated, the gels were washed twice with PBS and the PDMS stencils were carefully removed.

### Seeding on PAA substrates

3D intestinal organoids were mechanically disaggregated by pipetting in PBS with calcium and magnesium (Sigma-Aldrich). For one PAA gel, we seeded the number of organoids contained in one Matrigel drop of the 24 well plate. After disaggregation, the organoids were centrifuged at 100xg RT for 3.5 minutes and the pellet was resuspended in ENR medium. The organoids were seeded in a small volume (50µL) on top of the ECM-coated PAA gels and incubated at 37^°^C 5% CO_2_. After 1hour incubation, 550 µL of ENR medium was added on top of the previous 50µL. All experiments were performed 2-4 days after seeding.

### Immunostainings

The organoid monolayers were fixed in 4% Paraformaldehyde (Electron Microscopy Sciences) for 10 minutes at RT and washed three times with PBS. For Olfm4, GFP, Ki67, CK20 and Lysozyme, the samples were permeabilized with 0,1% Triton X-100 (Sigma-Aldrich) for 10 minutes at RT. After three washes with PBS, the samples were blocked with PBS containing 10% FBS (Gibco) for 1 hour at RT. Primary antibodies diluted in PBS containing 10% FBS were added and incubated overnight at 4^°^C. After three more washes in PBS, secondary antibodies in PBS containing 10% FBS were added for 1h at RT, washed 5 times with PBS (5 minutes each) and imaged.

For phalloidin, the samples were permeabilized with 0.5% Triton X-100 for 30 minutes at RT. After three washes with PBS, the samples were blocked with PBS containing 10% FBS and 0.2% Triton X-100 for 2 hours at RT. The samples were then incubated in PBS containing 10% FBS and 0.2% Triton X-100 overnight at 4^°^C. After 5 washes with PBS (5 minutes each wash), samples were incubated with phalloidin diluted in PBS containing 10% FBS for 1h at RT. Finally, the samples were washed 5 times with PBS (5 min each wash) and imaged.

### Antibodies

The primary antibodies used and their respective dilutions were: rabbit anti Olfm4 1:200 (Cell signaling Cat D6Y5A), mouse anti CK20 1:50 (Dako Cat M7019), mouse anti Ki67 1:100 (BD Biosciences Cat BD550609), rabbit anti Lysozyme 1:2000 (Dako Cat A0099), rabbit anti ZO-1 1:200 (Invitrogen Cat 402200) and mouse anti GFP 1:400 (Abcam, ab1218). The secondary antibodies used were: goat anti mouse Alexa Fluor 488 (Thermofisher Cat A-11029), Donkey anti rabbit Alexa Fluor 488 (Thermofisher Cat A-21206) and goat anti rabbit Alexa Fluor 555 (Thermofisher Cat A-21429). All secondary antibodies were used at 1:400 dilution. To label F-actin, Phalloidin Atto 488 (Sigma-Aldrich Cat 49409) was used at 1:500 and Phalloidin-iFluor 647 (Abcam Cat ab176759) was used at 1:400.

### Image acquisition

All images except laser cuts were acquired in a Nikon TiE inverted microscope with a spinning disk confocal unit (CSU-WD, Yokogawa) and a Zyla sCMOS camera (Andor). For the Supplementary Video 4, a 40× objective (Plan Fluor, NA 0.75, Dry) was used. For the rest of images, a 60× objective (Plan APO, NA 1.2, water immersion) was used. For life imaging experiments, a temperature box maintaining 37^°^C in the microscope (Life Imaging Services) and a chamber maintaining CO_2_ and humidity (Life Imaging services) were used. The open source Micromanager^55^ was used to carry out multidimensional acquisitions with a custom-made script.

To image the monolayers on 0.7kPa gels (Figs. 3,4), they were mounted upside down to improve image quality. Briefly, the samples were fixed in PFA 4% 10 minutes at RT. They were mounted in PBS and sealed with nail polish before being flipped and imaged from the apical side.

For laser ablation, images were acquired in a laser-scanning confocal microscope (LSM880, Carl Zeiss) at a resolution of 512×512 pixels (pixel size = 0.2595 µm) with a bidirectional scan. A Plan-Apochromat 40x objective (Plan Apochromat, NA 1.3, Oil, DIC M27) was used. The microscope was equipped with temperature control (37^°^) and CO_2_ and humidity control. Zeiss ZEN software was used to carry out the acquisitions.

### Three-dimensional traction microscopy

3D tractions were computed as previously described. Briefly, confocal stacks of the top layer of the fluorescent beads embedded in the polyacrylamide gels were imaged with a z-step of 0.2µm both in the deformed (by the cells) and relaxed (trypsinized) states. From the stacks, the 3D deformations of the gel were computed with a home-made iterative 3D Particle Image Velocimetry software^22^. A window size of 64×64 pixels and an overlap of 0.75 was used. A Finite Element Method solution was implemented to compute 3D tractions from the 3D substrate displacements^22^.

### Cell Velocities

Velocity of the monolayer was computed by performing Particle Image Velocimetry (PIV) analysis on consecutive time-lapse images of organoid monolayers expressing membrane-targeted tdTomato. To characterize cell kinematics in Fig. 5a, b, e, a window size of 64×64 pixels and an overlap of 0.75 was used.

### Radial averaging and kymographs

The boundary between the stem cell compartment and the transit amplifying zone was drawn by connecting the points where the normal traction component changed sign from negative to positive. If tractions were not available (as in immunostainings) the boundary was defined by the change in cell morphology (from columnar to elongated). For averaging (traction, velocity or image intensity), every pixel in the image was assigned the value of the distance to the closest point of the contour of the stem cell compartment. Spatiotemporal diagrams (kymographs) of a given variable were obtained by binning the values of that variable as a function of the (signed) distance from the boundary of the stem cell compartment and then computing the mean within each bin. For vectors, the normal to the wound contour was determined by fitting a parametric parabola to the 9 pixels surrounding each pixel of the contour. To attribute a normal vector to each pixel of the image we followed the approach of Trepat et al^35^. Briefly, we first assigned to each pixel of the image the normal vector of the closest pixel of the contour of the stem cell compartment. We then smoothed curvature and normal vector components using a moving average filter whose characteristic length increased with the distance from the contour. To build spatiotemporal kymographs, the process of masking and radial averaging was repeated for every single timepoint. Additional crypts in the field of view were excluded from the analysis.

In Fig. 3c, the normal traction of the stem cell compartment was computed as the mean of the normal traction at the crypt center. The normal traction of the transit amplifying zone was computed as the peak in positive normal traction within the first 20 microns outside the stem cell compartment.

### Averaging of crypt radial profiles

Crypts are variable in size within a certain range. As a consequence, directly averaging profiles of tractions, velocity or any other parameter of the study generates artifacts coming from the misalignment of compartments between different crypts. To avoid this, individual crypt profiles were linearly resized to the average crypt radius of the experimental group. Resized profiles were then averaged.

### Blebbistatin treatment

Organoid monolayers on 5kPa gels (Fig. 3a-c) were imaged for 1 hour 30 minutes in normal ENR medium (baseline). Then, ENR medium containing Blebbistatin (-/-) (Sigma-Aldrich) was added to a final concentration of 15µM and the monolayers were imaged for 3 hours. The medium was then aspirated, the sample was washed with ENR, and the monolayers were imaged for additional 11h in normal ENR.

Monolayers on 0.7kPa gels (Fig. 3d, e, Fig. 4b) or 15kPa gels (Fig. 4g, h) were treated with 15µM of Blebbistatin or DMSO (Sigma-Aldrich) in ENR medium for 3h. They were then fixed in 4% PFA for 10 minutes at room temperature. Finally, they were washed twice with PBS and imaged as indicated before. The monolayers of Extended Data Fig. 5 were imaged without fixation. Crypt indentation (Fig. 3e) was quantified in Fiji by tracing a vertical line from the bottom of the crypt to the bottom of the TA cells in a lateral view of the monolayer.

### Laser cuts

Laser ablation was performed using a Ti:Sapphire laser (Mai Tai DeepSee, Spectra Physics) set at 800 nm and laser power of 15% (500-750 mW). Circular regions for ablation were selected manually at the boundary between crypt compartments based on basal cell morphology. The cuts were performed in the medial plane of the monolayer.

To compute recoil velocities, PIV was performed using a window size of 16×16 pixels and an overlap of 0.75. Velocity fields were resized to the original image size (512×512) using a bicubic interpolation to compute velocities relative to the ablated region. A peak in tissue velocity occurred within the first 2 seconds after ablation and thus recoil velocity was computed in this timeframe.

For circular ablations (Fig. 3g, h), the contour of the ablated region was used to compute radial and tangential components of the recoil velocity as explained above. Radial velocities were averaged according to their distances to the ablated region. Crypt indentation before and after ablation was computed from the deformation of fluorescent beads embedded in the substrate as explained in the traction force microscopy section.

For ablations radial to the crypt (Fig. 5f), a straight line was fitted to the ablation region. The direction of this line was used to compute parallel and perpendicular components of the recoil velocity. Parallel velocity was considered positive when pointing away from the crypt and negative when pointing towards the crypt. Perpendicular velocity was considered positive when pointing away from the cut, and negative when pointing towards the cut. Quantification of recoil velocity was performed on bands of 50 pixels (12,975 mm) width at each side of the ablated region.

### Single cell shape analysis

#### Image pre-processing

Spinning disc images of intestinal organoid monolayers were pre-processed to address low signal-to-noise ratio and image scattering at apical and basal cell surfaces, presumably due to rich apical excretion, brush microvillar border, and scattering in the hydrogel underlying the organoid monolayers. To enable automatic detection of all single cell surfaces, signal intensity values in dim regions of apical and basal surfaces were computationally increased. Using Fiji^56^, monolayer apical and basal surface was defined in 3D, following the contour of the monolayer by manually adding sparse points in the xz-view and interpolating the surface between them, using the Volume Manager plugin (by Robert Haase, MPI-CBG). Using a custom Matlab script, image pixels lying on the capping surface and having fluorescence signal intensity higher than 1st and lower than 3rd quartile of overall image intensity were set to the 3rd quartile value (4th quartile in case of extremely bright lateral cell surfaces), resulting in segmentable apical and basal membrane signal in cells that displayed only clear lateral surfaces in original images.

#### Cell segmentation

Pre-processed cell membrane images were used for automatic cell segmentation in MorphographX^57^. Images were processed with Gaussian blurring (sigma 0.3-0.5) and stack normalization (radius 4-10, blur factor 0.5-0.8). Segmentation was performed in whole image stacks in 3D using the Insight Toolkit watershed algorithm implemented in MorphographX (threshold 800-1500). All segmentations required extensive manual correction for oversegmented, undersegmented and incomplete cells. 3D cell meshes were created (cube size 2, smoothing 1) from pixel label images for further shape analysis.

#### Shape analysis

Shape analyses were performed in 3D using custom-developed Matlab tools. To account for the complex shape of cells in the organoid monolayers, an apicobasal cell axis was constructed for each cell from centroids of apicobasal cell cross-sections. This 3D axis was used to calculate 3D cell height and to obtain cross-sections normal to the curved apicobasal axis. Obtained sections were analyzed for their area and various shape parameters, resulting in detailed apicobasal profiles for all cells. Cell tilt to crypt center was defined as the angle of the straight, end-to-end cell axis with respect to the z-axis, and set to be <0 if the axis was pointing away from the crypt center. Cell position was calculated as the 2D Euclidean distance of the basal cell centroid from crypt center, defined as a center of crypt mask manually outlined in the basal-most tissue plane in Fiji. Paneth cells were manually identified in 3D segmentations based on their distinct (granular) apical appearance in corresponding bright field images.

### F-Actin and myosin quantification

Given the height heterogeneity between compartments in the organoid monolayer, we developed the following algorithm to define the cell apical, medial and basal coordinates for every XY position of the substrate. For F-actin stainings (phalloidin), each plane of the imaged Z-stack was first binned (8×8). Then, for each XY position of the binned image along the substrate, the apical and basal coordinates were defined as the coordinates of the apical-most and basal-most peaks in fluorescence intensity (phallodin). The medial position was defined as the Z coordinate equidistant to the apical and basal coordinate for each XY pixel. We then extracted the fluorescence intensity of the apical, medial and basal positions. A similar approach was used to quantify myosin IIA, but using the Volume Manager plugin (by Robert Haase, MPI-CBG) to segment the monolayer.

### Quantification of cell density

Maps of the spatial distribution of cells were manually obtained using Fiji multipoint selections. The coordinates of each cell were imported to Matlab. The mask of the stem cell compartment was used to compute the radial distance to the crypt center of each pixel of the field of view as explained above. Radial coordinates were divided in bins of 5 mm width to quantify the number of cells per unit area and generate radial density profiles.

### Graphs and statistics

Traction maps, kymographs and laser cuts were plotted in Matlab. The rest of the plots were generated in Graphpad Prism 8. Immunostainings and videos were processed with the open source sofwares Fiji and HandBrake. The data is represented as mean ± error (SEM or SD as indicated in the figure captions). Normality of the data was checked and the statistical test used to compare means was chosen accordingly. All the statistical analyses were performed in Graphpad Prism 8.

## Code availability

MATLAB analysis procedures are available from the corresponding authors on reasonable request

## Data availability

The data that support the findings of this study are available from the corresponding authors on reasonable request

## Acknowledgements

We thank Jorge Barbazán for assistance with laser ablations and discussions; Ernest Latorre for support with image segmentation and traction software; Fatima el Marjou, Andrew Clark, Myriam Ali Ziane, Carme Cortina and Xavier Hernando for support and training on organoid culture and *in vivo* procedures; and Ariadna MarÍn, Nimesh Ramesh Chahare, Tom Golde, Juanfra Abenza, Adam Ouzeri and all members of the Roca-Cusachs, Vignjevic, Arroyo and Trepat labs for discussions and support. Funding: the authors are funded by Spanish Ministry for Science, Innovation and Universities MICCINN/FEDER (PGC2018-099645-B-I00 to X.T., DPI2015-71789-R to M.A.), the Generalitat de Catalunya (Agaur, SGR-2017-01602 to X.T., 2014-SGR-1471 to M.A., SGR-2017-01602 to E.B. the CERCA Programme, and “ICREA Academia” award to M.A.), the European Research Council (CoG-616480 to X.T., CoG-681434 to M.A., CoG-772487 to D.M.V, AdvG 340176 to E.B.), the European Union’s Horizon 2020 research and innovation programme under the Marie Skłodowska-Curie grant agreement No 797621 to M.G-G. and No. 792028 to F.G., Obra Social “La Caixa” (ID 100010434, fellowship LCF/BQ/DR19/11740013 to G.C., LCF/BQ/DE14/10320008 to C.P-G. and 01/16/FLC to A.A-V.), Fundació la Marató de TV3 (project 201903-30-31-32 to XT and EB). IBEC, IRB and CIMNE are recipients of a Severo Ochoa Award of Excellence from the MINECO.

## Author contributions

C.P-G. and G.C. performed all experiments except for immuno-stainings, which were also performed by N.C. C.P-G., G.C., M.M. and M.G-G. developed analysis software and analyzed data. M.M. performed image segmentation. F.G., S.K. and M.A implemented the computational model. A.A-V., P.R-C, E.B. contributed technical expertise, materials, and discussion. C.P-G, D.M.V., M.A., and X.T. conceived the project. D.M.V., M.A., and X.T. supervised the project. C.P-G, G.C., M.A., and X.T. wrote the manuscript.

## Additional information

**Extended Data** is available for this paper

**Supplementary Information** is available for this paper

**Correspondence and requests for materials** should be addressed to X.T., M.A. or D.M.V.

