## Supplementary information for "Mechanical compartmentalization of the intestinal organoid enables crypt folding and collective cell migration"

### Correspondence to:

### Extended Data

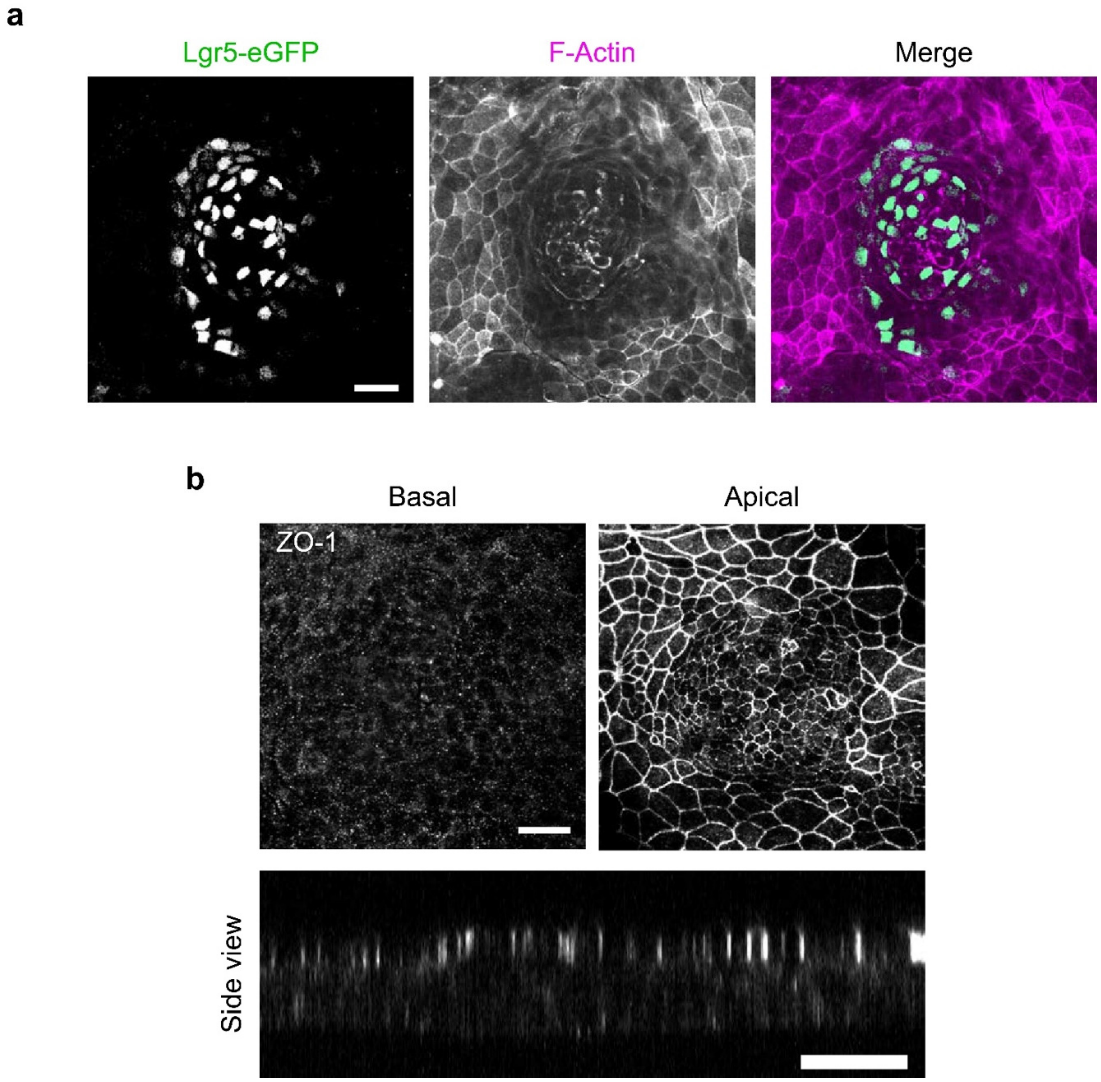


**Extended Data Fig. 1 | Compartmentalization of stem cells and apicobasal polarity in organoid monolayers.**
**a,** Organoid monolayers expressing Lgr5-eGFP-IRES-CreERT2 stained for GFP and F-actin (phalloidin). Scale bar, 20 µm. Stiffness of the gel, 5kPa. **b,** Organoid monolayers stained for Zonula occludens 1 (ZO-1). Top: Average intensity projections of the 10 most basal (Top left) or apical (Top right) planes of the monolayer. Bottom: lateral view of the monolayer. Scale bars, 20 µm. Stiffness of the gel, 5kPa.

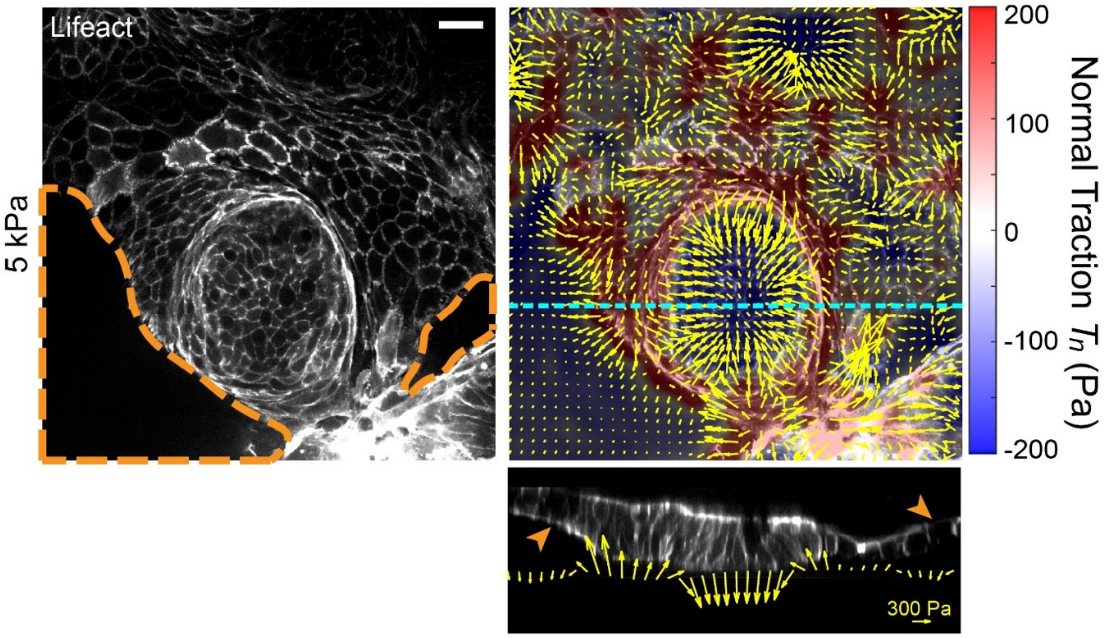


**Extended Data Fig. 2 | Spontaneous formation of pressurized domes in the villus-like domain.**

Top left: medial view of an organoid monolayer expressing Lifeact-eGFP on a 5kPa gel. The dashed orange line defines the regions where the monolayer has delaminated to form a pressurized dome (orange arrowheads in bottom panel). Top right: 3D traction map of the same crypt. Yellow vectors represent components tangential to the substrate and the color map represents the component normal to the substrate. Horizontal cyan line indicates y-axis position of the lateral *xz* view (bottom). Bottom: lateral view of the organoid monolayers. Yellow vectors represent tractions. Scale bar, 20 µm. Scale vector, 300Pa.

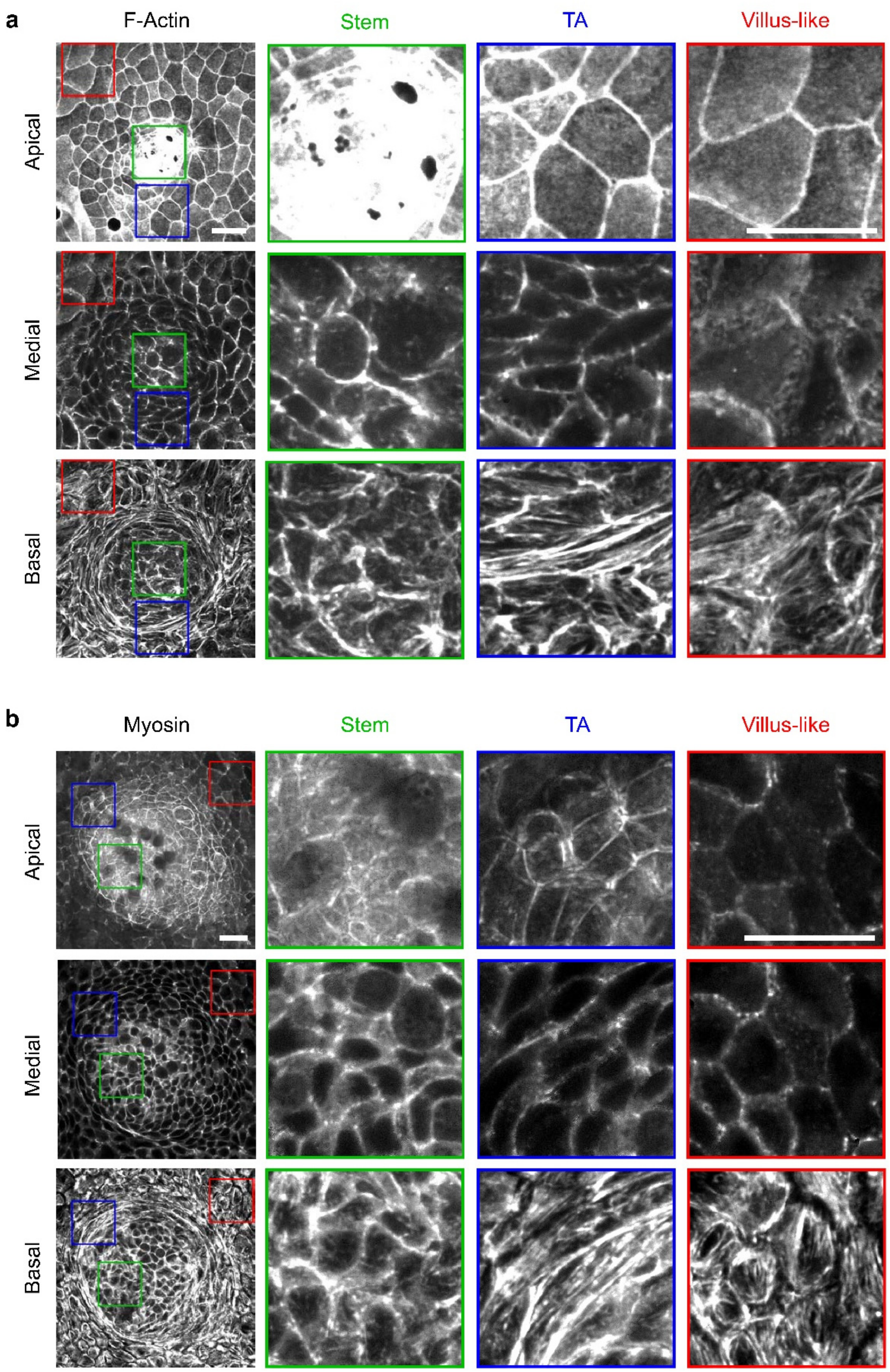


**Extended Data Fig. 3 | Apicobasal distribution of F-actin and Myosin in organoid monolayers.**

**a-b,** Apical, medial and basal projections of F-Actin (Phalloidin, **a**) and myosin IIA-eGFP (**b**). The stem cell compartment (Stem), the Transit amplifying zone (TA) and the villus-like domain (villus-like) are zoomed in the regions of the monolayer indicated with the respective colors. Scale bars, 20 µm. Stiffness of the gel, 5kPa.

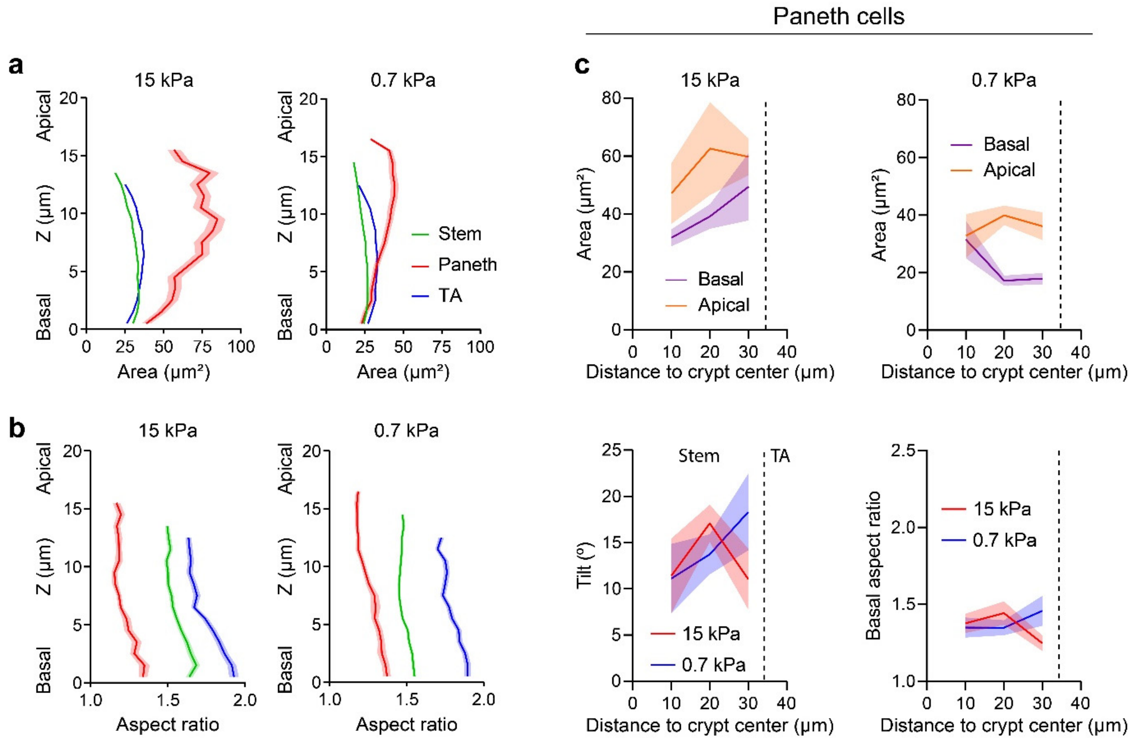


**Extended Data Fig. 4 | Morphometric analysis of the different cell types in the crypt.** **a-b,** Cell area (**a**) and aspect ratio (**b**) along the apicobasal axis of stem (green), Paneth (red) and TA (blue) cells on rigid (left, 15kPa) and soft (right, 0,7kPa) gels. N = 262 (stem cells), N = 20 (Paneth cells) ; N = 200 (transit amplifying cells) for 15kPa gels. N = 614 (stem cells); N = 51 (Paneth cells); N = 307 (transit amplifying cells) for 0,7kPa gels. N = 3 independent crypts per stiffness from 2 (0,7kPa) and 3 (15kPa) independent experiments. Data are represented as mean ± SEM. **c**, Top: Apical and basal area of Paneth cells as a function of the distance to the crypt center on stiff (left, 15 kPa) and soft (right, 0.7 kPa) substrates. The boundary between the stem cell compartment and the transit amplifying zone is indicated in all the plots with a dashed vertical line. Bottom: Apicobasal tilt (left) and basal aspect ratio (right) of Paneth cells as a function of the distance to the crypt center on stiff (red, 15 kPa) and soft (blue, 0.7 kPa) substrates. N = 3 independent crypts per stiffness from 2 (0,7kPa) and 3 (15kPa) independent experiments. From center to edge bins, N = 6, 6 and 8 cells for 15 kPa gels and N =11, 25 and 15 cells for 0,7kPa gels. Data are represented as mean ± SEM.

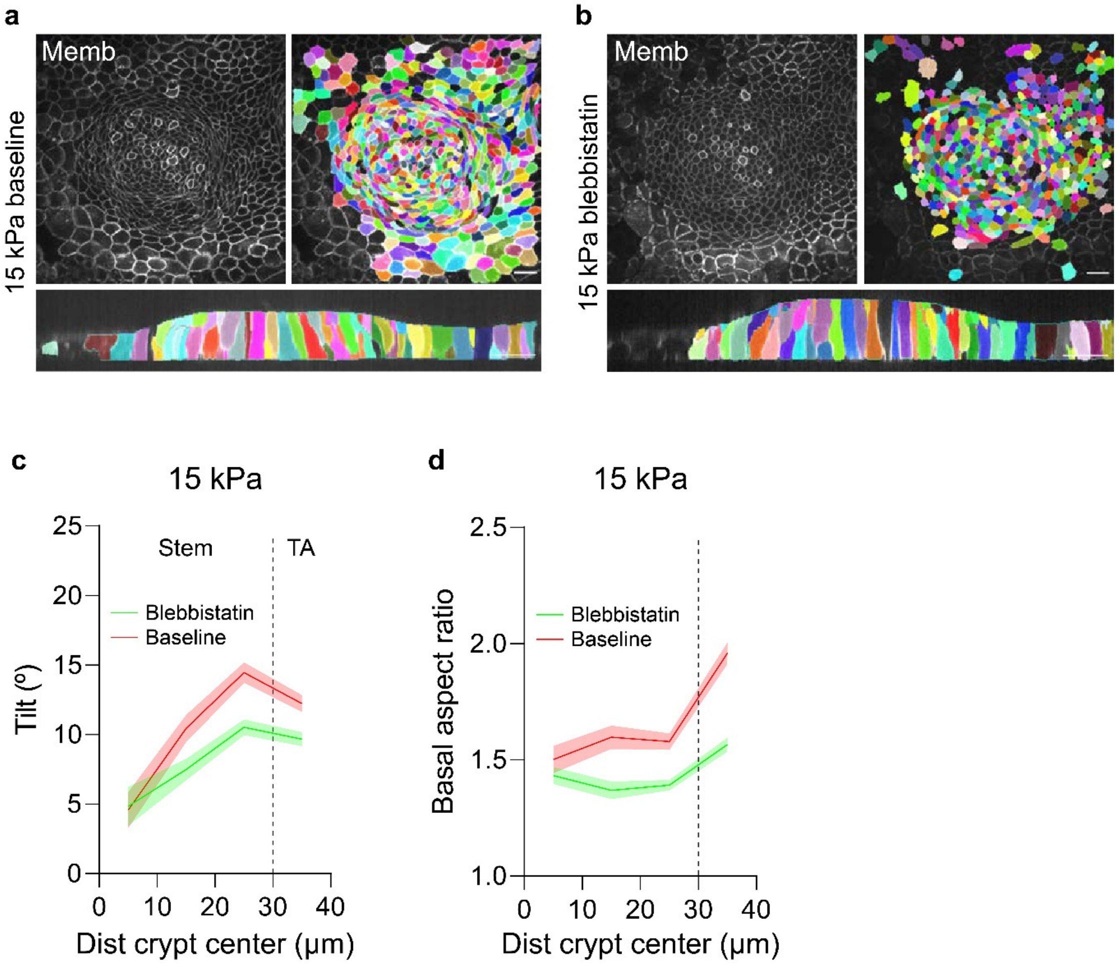


**Extended Data Fig. 5 | Effect of myosin inhibition in organoid cell shape.**
**a-b,** 3D segmentation of a crypt on 15 kPa gels under baseline conditions (**a**) and the same crypt after 3h treatment with 15µM of blebbistatin (**b**). Top: medial view. Bottom: lateral view. Scale bar, 20 µm. **c-d,** Apicobasal tilt (**c**) and basal aspect ratio (**d**) as a function of the distance to crypt center on rigid substrates (15kPa) before and after blebbistatin. Vertical dashed line indicates the boundary between the stem cell compartment and the transit amplifying zone. From center to edge bins, N=42, 71, 146 and 204 cells for baseline crypt and 39, 81, 126 and 92 for blebbistatin treatment. N= 3 independent crypts per condition from 2 independent experiments.

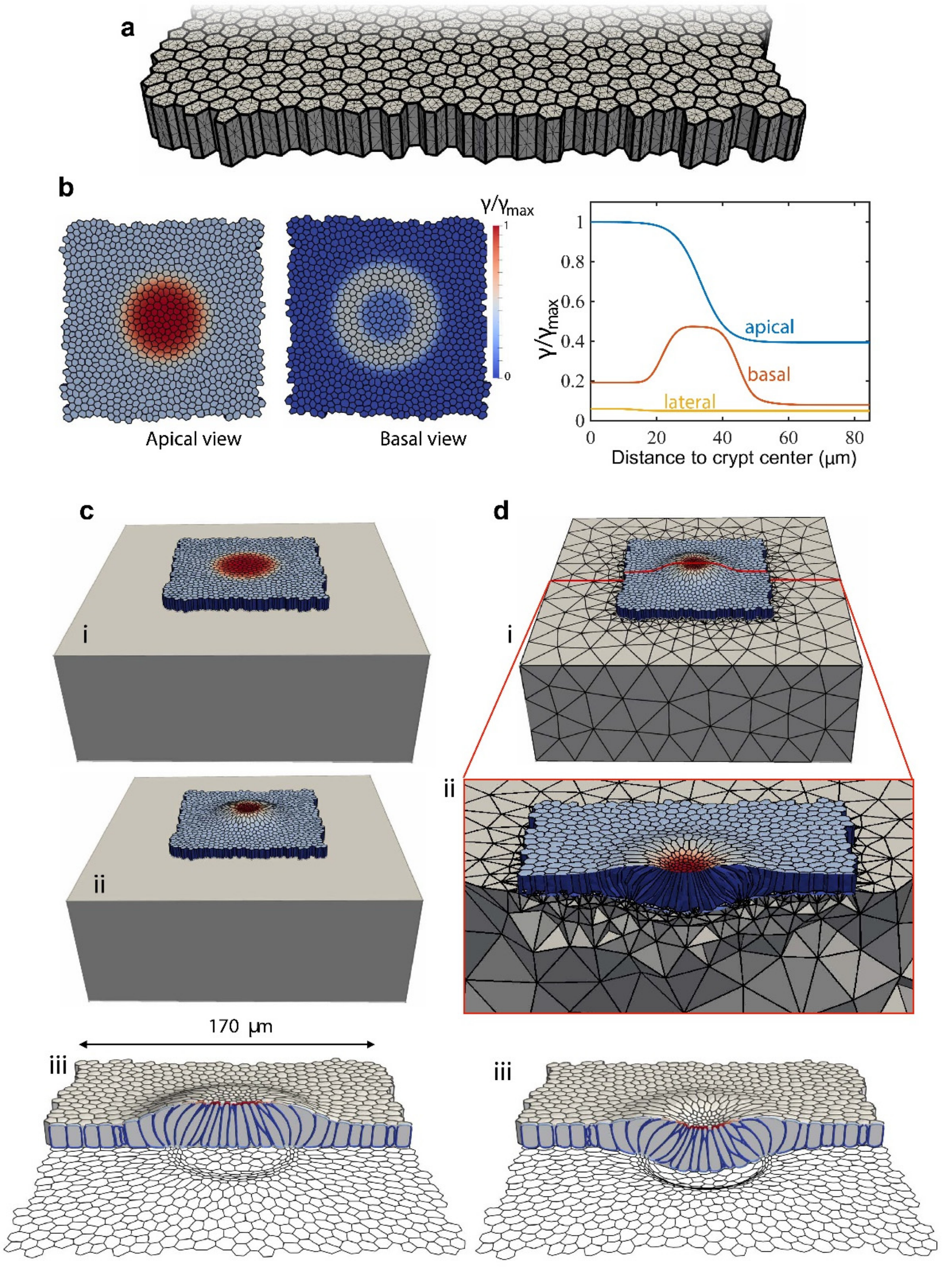


**Extended Data Fig. 6 | 3D computational vertex model and simulation protocol.**
**a**, Discretization of the tissue: the thick lines denote the intersection between cellular faces and the thin lines the triangulation of the cell surfaces. **b**, Pattern of apical, basal and lateral surface tensions prescribed in the initial regular cell monolayer. **c**, Equilibration of the initial regular monolayer with patterned surface tensions on a rigid substrate, where basal nodes are constrained to a plane but can slide horizontally. Initial state (i), equilibrated state (ii), and different view of equilibrated state with basal cell outline (iii). **d**, Coupling with a deformable substrate, modeled computationally with a tetrahedral mesh discretizing a hyperelastic block (i). The equilibrated crypt on a rigid substrate (**c**-ii) is further equilibrated on the deformable substrate (**d**-ii,iii).

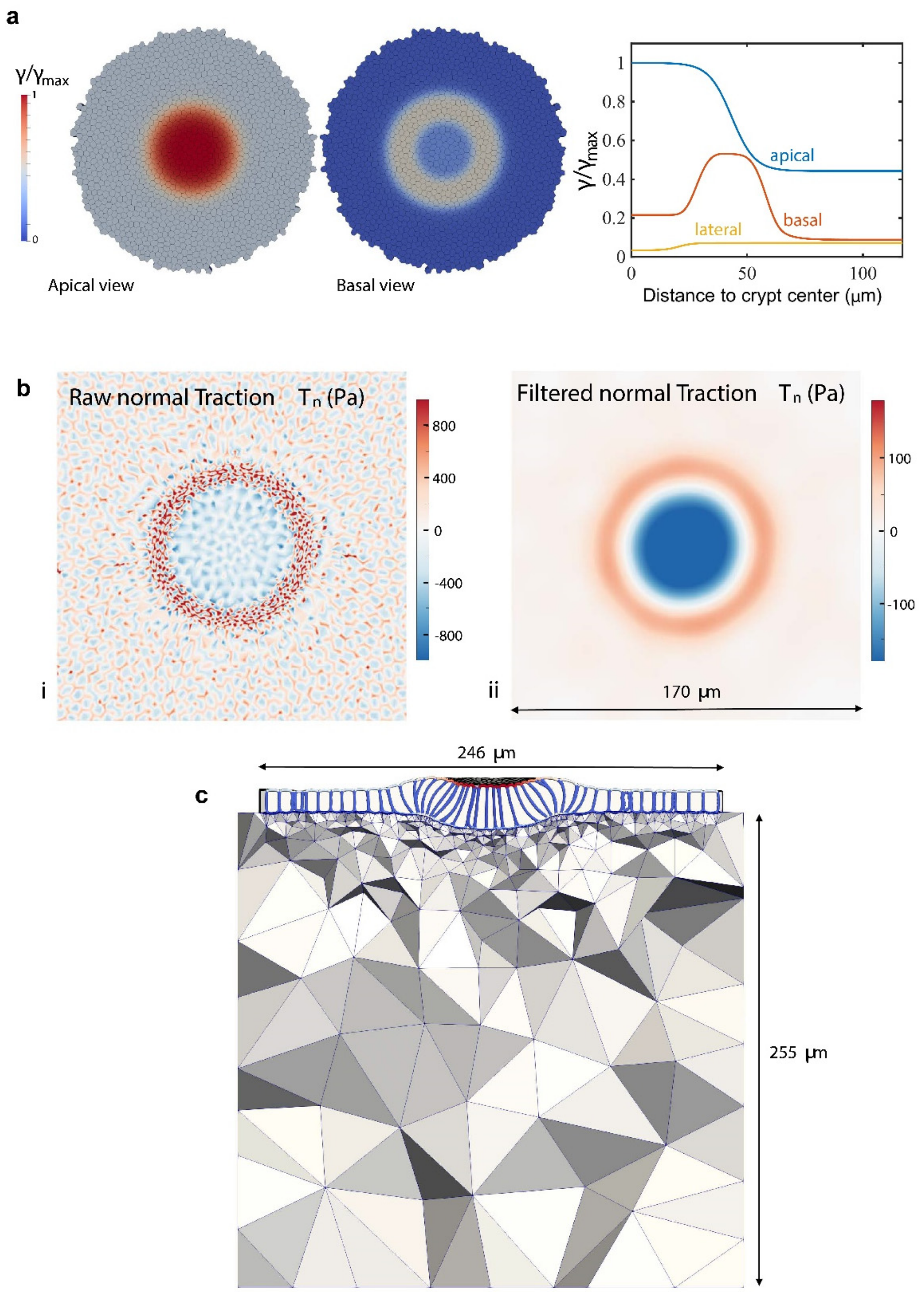


**Extended Data Fig. 7 | Representative crypt shown in Fig. 4.**
**a**, Pattern of apical, basal and lateral surface tensions prescribed in the initial regular cell monolayer. **b**, Maps of basal normal traction. (i) Raw normal tractions at the basal plane featuring sub-cellular fine-scale details. To compare with experimental averages, we filtered these tractions with a Gaussian filter with standard deviation of 6 µm, (ii). **c**, Computational model of the deformed crypt on a soft hyperelastic substrate.

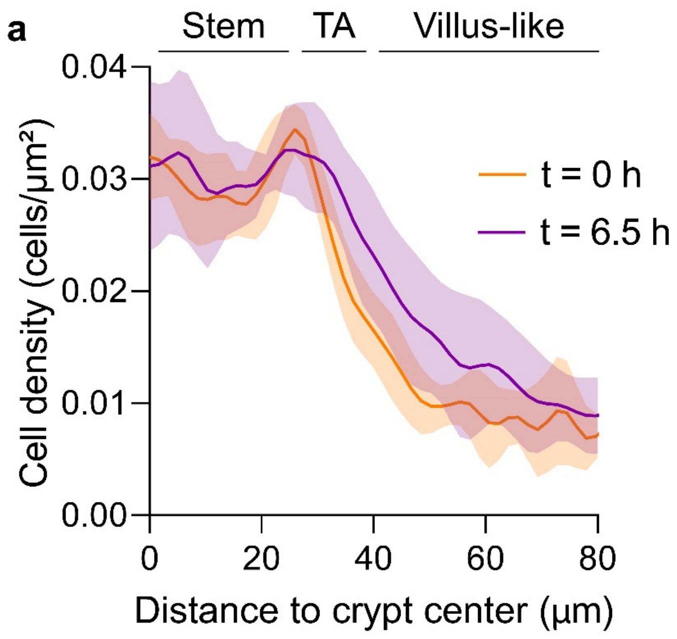


**Extended Data Fig. 8 | Cell density profile in the organoid monolayers.**
**a**, Cell density as a function of the distance to the crypt center at the beginning (t=0h, orange curve) and at the end (t=6.5h, purple curve) of the experiment. Data are represented as mean ± SD of the 5 crypts in Fig. 5,b-e .

### Supplementary note 1: Computational vertex model.

To understand the tissue mechanics leading to cell and monolayer morphology and to the mechanical coupling with the substrate, we developed a 3D computational vertex model. This model is based on a conventional effective energy or virtual work function of the form

$$\delta W = \sum_{c=1}^{N} \sum_{f=1}^{N_c} \gamma_{f,c} \delta A_{f,c}, \tag{1}$$

where $N$ is the number of cells, $N_c$ the number of faces of cell $c$, $\gamma_{f,c}$ the surface tension of face $f$ of cell $c$, and $\delta A_{f,c}$ the variation of the surface area of that face. We assume that the surface tensions $\gamma_{f,c}$ remain constant during a simulation but are heterogeneous throughout the tissue. Work functionals for 3D vertex models can also account for the line tension generated by apical or basal cables. We also implemented such terms but found no essential differences for the purpose of this study, and hence we ignored them for the sake of simplicity.

To capture the cell shapes with curved junctions observed in the experiments, we discretized each cell with a triangulation as shown in Extended Data Figure 6a. Accounting for cell volume preservation, we can define an effective or pseudo-energy over the entire triangulation describing the tissue of the form[28–30]

$$W(\boldsymbol{x}_1, \ldots, \boldsymbol{x}_M) = \sum_{c=1}^{N} \sum_{f=1}^{N_c} \gamma_{f,c} A_{f,c}(\boldsymbol{x}_1, \ldots, \boldsymbol{x}_M) + \sum_{c=1}^{N} \frac{\kappa}{2} \left[ V_c(\boldsymbol{x}_1, \ldots, \boldsymbol{x}_M) - V_{c,0} \right]^2 \tag{2}$$

where $\boldsymbol{x}_i$ denotes the position of node $i$ in this triangulation, $V_c$ is the volume of cell $c$, and $\kappa$ is an osmotic compressibility modulus that ensures that cell volumes remain fixed within 0.1% to the initial volume, $V_{c,0}$ for cell $c$.

We started our analysis by considering a planar tissue with cells of uniform shape and size as shown in Extended Data Figure 6. To examine the mechanics of intestinal crypts, we prescribed a distribution of surface tensions that mimics the measured basal and apical F-actin distribution. It is known that cortical tension depends in a highly nontrivial way on the amount, but also the architecture, of cytoskeletal components. However, given the order-of-magnitude variations of F-actin accumulation in our crypts, it is reasonable to consider

F-actin accumulation as a proxy for cortical tension in a first approximation. Noting that the F-actin distributions were measured on an actual deformed crypt and that we are prescribing surface tensions on an idealized undeformed tissue, we broadened the apical and basal peaks in F-actin distribution in our computer model. Regarding lateral surface tensions, we noted that the apical/basal surfaces of our monolayers were quite smooth at the intersections with lateral junctions. We reasoned that if lateral surface tensions were relatively large, then as a result of mechanical equilibrium at these intersections we should observe noticeable surface deformations at apico-lateral and baso-lateral intersections. Since these were absent, we concluded that lateral tensions should be significantly smaller than basal and apical tensions. We note in this regard that at lateral faces, adhesion tension acts as a negative surface tension that lowers the total lateral surface tension.

During our analysis, we kept fixed the distribution of surface tension over cell faces shown in Extended Data Figure 6b. Given the high heterogeneity of this surface tension pattern, the initial regular cell monolayer is not in mechanical equilibrium. We then proceeded to the equilibration of the system on a rigid substrate. For this, we minimized the function in Eq. (2) using Newton's method combined with a line-search algorithm to find the equilibrium positions of nodes in our triangulation. While our description of each cell face as a triangulated surface allows us to describe curved shapes, it also poses a numerical challenge as the distortion of our triangulations needs to be controlled during the numerical minimization. Indeed, since the function in Eq. (2) only depends on the surface area of each face and the volume of each cell, it is invariant with respect to tangential motions of internal nodes to each face that leave these geometric quantities unchanged, and thus does not provide any control of mesh distortion. To deal with this issue, we adopted the approach proposed elsewhere[59], which considers a fictitious surface hyperelastic model for each cell junction whose reference configuration is updated iteratively to the previously converged configuration. This fictitious elastic energy controls mesh distortions and upon convergence of the algorithm it vanishes and does not bias the final results.

In our simulations on rigid substrates, we allowed the nodes on the basal plane to slide tangentially but forced their $z$ position to zero so that they stay on the plane of the substrate. Furthermore, we fixed the lateral edges of the tissue. Upon equilibration, we found the tissue and cellular shapes and the normal tractions, Extended Data Figure 6c. As the tissue deformed, the pattern of surface tensions more closely followed the experimentally measured patterns of F-actin distribution.

We then placed the equilibrated crypt in contact with a highly deformable elastic substrate. We modeled the substrate using finite deformation continuum mechanics and a NeoHookean hyperelastic model, for which the strain energy density per unit undeformed volume is given by[60]

$$\psi(\boldsymbol{C}) = \frac{\lambda}{2}(\ln J)^2 - \mu \ln J + \frac{\mu}{2}(\text{trace}\,\boldsymbol{C} - 3)\,, \tag{3}$$

where $\boldsymbol{C} = \boldsymbol{F}^T\boldsymbol{F}$ is the right Cauchy-Green deformation tensor, $\boldsymbol{F}$ is the deformation gradient, $J = \sqrt{\det \boldsymbol{C}}$ is the Jacobian determinant and $\lambda$ and $\mu$ are the Lamé coefficients at infinitesimal deformations. These coefficients are related to infinitessimal Young's modulus

$E$ and Poisson's ratio $\nu$ by the relations $\lambda = E\nu/[(1+\nu)(1-2\nu)]$ and $\mu = E/(1+\nu)$. We discretized the deformable substrate with linear tetrahedral finite elements. The bulk tetrahedral mesh was generated to be conforming in its top plane to the surface triangulation of the basal plane of the tissue equilibrated on a rigid substrate. We imposed kinematical compatibility at the cell-matrix interface by identifying the basal nodes of the tissue triangulation to the corresponding nodes of the matrix mesh. We then further equilibrated the joint tissue-matrix system by minimizing the joint effective energy given by

$$W + \int_{\Omega_0} \psi(\boldsymbol{C})\, dV, \tag{4}$$

where $\Omega_0$ is the domain representing the substrate, with respect to the positions of the nodes of the tissue triangulation and of the bulk finite element mesh. This minimization was again performed using Newton's method with a line-search. As a result, the crypt was able to deform the substrate, Extended Data Figure 6d.

We simulated tens of computational crypts with patterns of cellular surface tensions following the measured F-actin distribution. We found a very robust agreement with the main features of the experiments in terms of tissue shape in stiff and soft substrate, of cellular shapes and of normal traction. However, the details of these observables slightly depended on the specific pattern of surface tensions. In Extended Data Figure 7a we report the pattern of surface tensions used to produce the results in Fig. 4. We also illustrate the procedure to filter the normal tractions in the simulations, Extended Data Figure 7b, and the finite element mesh used to model the substrate, Extended Data Figure 7c.

The length-scale of the model is set by the typical size of cells and height of the typical crypt. The force-scale of this model is given by $\gamma_{\max}$, Extended Data Figures 6b and 7a. By scaling the magnitude of the computationally obtained normal tractions to the measured ones, we could thus quantify the magnitude of the surface tensions predicted by the model. We found $\gamma_{\max} \sim 4.6$ mN/m, which is about two times larger than surface tensions measured in suspended cells during mitosis[58]. We note that apical actin cables could contribute to this effective apical surface tension.

### Supplementary note 2: Balance of cell number.

Let $\rho$ denote the number areal density of cells and $\boldsymbol{v}$ the velocity of cells. Balance of cell number in steady state can be written as

$$s = \text{div}\ (\rho \boldsymbol{v}), \tag{5}$$

where $s$ is the rate of cell addition per unit area. To evaluate $s$ from measurements, we measured and radially averaged $\rho(r)$ and $v(r)$ where $v$ is the radial velocity. Under axisymmetry, the equation above can be expressed as

$$s = \frac{d}{dr}(\rho v) + \frac{\rho v}{r}, \tag{6}$$

which we evaluated by fitting a spline to $\rho(r)v(r)$ and differentiating the spline.

### Supplementary note 3: Axisymmetric Monolayer Stress Microscopy.

We describe here a simple approach for Monolayer Stress Microscopy (MSM), that is to infer the tissue surface stress from the measured tractions, in an axisymmetric configuration pertinent to the analysis of our crypts. The starting variable is then the radially-averaged tangential traction $T_r$, Fig. 1f.

#### Background

We consider polar coordinates given by $x(r,\theta) = r\cos\theta$ and $y(r,\theta) = r\sin\theta$. The natural basis is given by $\boldsymbol{x}_r(r,\theta) = (\cos\theta, \sin\theta)$ and $\boldsymbol{x}_\theta(r,\theta) = (-r\sin\theta, r\cos\theta)$. Hence the metric tensor is given by

$$\{g_{ab}\} = \begin{pmatrix} 1 & 0 \\ 0 & r^2 \end{pmatrix}, \qquad \{g^{ab}\} = \begin{pmatrix} 1 & 0 \\ 0 & 1/r^2 \end{pmatrix}.$$

The corresponding Christoffel symbols are

$$\{\Gamma^r_{ab}\} = \begin{pmatrix} 0 & 0 \\ 0 & -r \end{pmatrix}, \qquad \{\Gamma^\theta_{ab}\} = \begin{pmatrix} 0 & 1/r \\ 1/r & 0 \end{pmatrix}.$$

Therefore, using standard formulae in differential geometry, we can compute the covariant derivative of a radial vector field $\boldsymbol{u}(r,\theta) = u(r)\,\boldsymbol{x}_r(r,\theta)$ as

$$\{u^a{}_{|b}\} = \begin{pmatrix} u' & 0 \\ 0 & u/r \end{pmatrix}.$$

The symmetrized displacement gradient is the small-strain tensor, which thus takes the form

$$\{\varepsilon_{ab}\} = \begin{pmatrix} u' & 0 \\ 0 & ur \end{pmatrix}, \qquad \{\varepsilon^{ab}\} = \begin{pmatrix} u' & 0 \\ 0 & u/r^3 \end{pmatrix},$$

and its trace is

$$\text{tr}\ \boldsymbol{\varepsilon} = \varepsilon^a{}_a = u' + u/r.$$

Consider a stress tensor in axisymmetry, and thus of the form

$$\{\sigma^{ab}\} = \begin{pmatrix} \sigma^{rr}(r) & 0 \\ 0 & \sigma^{\theta\theta}(r) \end{pmatrix}.$$

The radial component of its divergence $\sigma^{ab}{}_{|b}$ can be written as

$$\sigma^{rb}{}_{|b} = \partial_r \sigma^{rr} + \frac{1}{r}\left(\sigma^{rr} - r^2\sigma^{\theta\theta}\right) = \partial_r \sigma^r{}_r + \frac{1}{r}\left(\sigma^r{}_r - \sigma^\theta{}_\theta\right) \tag{7}$$

Consider now linear elastic constitutive relation

$$\boldsymbol{\sigma} = 2\mu\boldsymbol{\varepsilon} + \lambda(\text{tr}\ \boldsymbol{\varepsilon})\boldsymbol{g}.$$

Using the equations above, we find

$$\sigma^r{}_r = \sigma^{rr} = (2\mu + \lambda)u' + \lambda\frac{u}{r}, \tag{8}$$

and

$$\sigma^\theta{}_\theta = r^2\sigma^{\theta\theta} = (2\mu + \lambda)\frac{u}{r} + \lambda u', \tag{9}$$

and we obtain its radial divergence as

$$(\text{div}\ \boldsymbol{\sigma})_r = \left[(2\mu + \lambda)u' + \lambda\frac{u}{r}\right]' + \frac{2\mu}{r}\left(u' - \frac{u}{r}\right).$$

If material properties are uniform, then we obtain

$$(\text{div}\ \boldsymbol{\sigma})_r = (2\mu + \lambda)\left(u'' + \frac{u'}{r} - \frac{u}{r^2}\right).$$

### Monolayer Stress Microscopy

The tissue modeled as a 2D continuous medium is initially in an axisymmetric state of mechanical equilibrium characterized by a stress $\boldsymbol{\sigma}$. The equilibrium condition can be expressed as

$$(\text{div}\ \boldsymbol{\sigma})_r = \partial_r \sigma^r{}_r + \frac{1}{r}\left(\sigma^r{}_r - \sigma^\theta{}_\theta\right) = T_r, \tag{10}$$

where $T_r$ are the measured radial tractions exerted by cells on the substrate. Thus, we have one equation for two unknowns, $\sigma^r{}_r$ and $\sigma^\theta{}_\theta$. One standard way to proceed is then to assume a constitutive relation, e.g. linear elasticity[34], leading to

$$\left[(2\mu + \lambda)u' + \lambda\frac{u}{r}\right]' + \frac{2\mu}{r}\left(u' - \frac{u}{r}\right) = T_r \tag{11}$$

or if mechanical properties are assumed to be constant to

$$(2\mu + \lambda)\left(u'' + \frac{u'}{r} - \frac{u}{r^2}\right) = T_r, \tag{12}$$

which, along with boundary conditions $u(0) = 0$ and $u(r^\infty) = U$ allow us to find the auxiliary displacement $u(r)$. The boundary condition at a far away location $u(r^\infty) = U$ only fixes the undetermined hydrostatic and uniform state of tension $\sigma_0$, and is thus arbitrary unless an independent measurement is available. Finally, recalling Eqs. (8,9) we obtain the sought-after tension as

$$\sigma^r{}_r = \sigma_0 + (2\mu + \lambda)u' + \lambda\frac{u}{r}, \tag{13}$$

$$\sigma^\theta{}_\theta = \sigma_0 + (2\mu + \lambda)\frac{u}{r} + \lambda u'. \tag{14}$$

The quantity $\sigma^r{}_r$ is the radial tension reported in Fig. 5, whereas $\sigma^\theta{}_\theta$ is a hoop tension.

We lack an independent measurement to determine $\sigma_0$ but our laser cuts indicate that radial tension is positive where it is minimum, at the transit amplifying zone. Therefore, in our calculations of MSM reported in Fig. 5 we chose $\sigma_0$ so that the minimum radial tension is zero with the understanding that there should be a positive offset to this curve.

Given $T_r$, we solve Eq. (11) using 1D finite elements. For this, we multiply Eq. (11) by a test function $w$, integrate over the domain $(0, R_{\text{out}})$ noting that $dS = 2\pi r dr$, and integrate by parts, to obtain

$$\int_0^{R_{\text{out}}} (2\mu + \lambda)u'w'rdr + \int_0^{R_{\text{out}}} \frac{2\mu + \lambda}{r} uwdr + \int_0^{R_{\text{out}}} \lambda\left(vw' + v'w\right)dr = -\int_0^{R_{\text{out}}} T_r wrdr.$$

Approximating the unknown with finite element basis function $u(r) = \sum_{J=1}^{M} N(r)u_J$ and taking the test functions to be $w(r) = N_I(r)$, we find the following algebraic system of equations

$$\sum_{J=1}^{M} K_{IJ}u_J = F_I,$$

imposing the constraints that $u_1 = 0$ and $u_M = U$ and where

$$K_{IJ} = \int_0^{R_{\text{out}}} (2\mu + \lambda)N_I'N_J'rdr + \int_0^{R_{\text{out}}} \frac{2\mu + \lambda}{r} N_I N_J dr + \int_0^{R_{\text{out}}} \lambda\left(N_I N_J' + N_I' N_J\right)dr$$

and

$$F_I = \int_0^{R_{\text{out}}} T_r N_I rdr.$$

In the calculations reported in Fig. 5, we chose $\lambda = 10\mu$. We checked that the inferred tensions were largely insensitive to the choice of material parameter provided that $\lambda \gtrsim 3\mu$. Since the crypt is bound to have very different material properties than the rest of the tissue, we tested the sensitivity of the recovered tensions to heterogeneous elastic constants. For this, we chose $\lambda$ and $\mu$ to be 10 and 20 times larger in the crypt than in the rest of the tissue. The results were again insensitive.

### Supplementary Videos

**Supplementary Video 1. Spreading of intestinal crypts**
Spreading of intestinal crypts on 5kPa polyacrylamide gels coated with collagen I and laminin 1. Over the course of three days, a 2D monolayer with crypt-like and villus-like domains is established. Membrane is labelled with tdTomato. Total duration: 45 hours.

**Supplementary Video 2. Crypt dynamics.**
Time-lapse of a basal (left) and apical (right) plane of the organoid monolayer. Multiple cell divisions can be observed at the crypt (center). Cells migrate collectively out of the crypt. Membrane is labelled with tdTomato. Total duration: 6 hours 32 minutes.

**Supplementary Video 3. Tracking of single cells in their journey out of the crypt.**
One cell (yellow) starts at the stem cell compartment and enters into the transit amplifying zone, acquiring an elongated shape. Another cell (light blue) at the transit amplifying zone divides and enters in the villus-like domain. Some cells (dark blues, light blue and green) enter in the villus-like domain and accelerate, separating from their followers. Membrane is labelled with tdTomato. Total duration: 6 hours 32 minutes.

**Supplementary Video 4. Division, migration, extrusion and doming in organoid monolayers.**
Time-lapse of a medial (left) and apical (right) plane of an organoid monolayer. A crypt in the center exhibits multiple cell divisions and radial migration into the villus-like domain. Multiple epithelial domes appear at times 1.20 - 2h (top right) and 8.40 h (left). Cells are constantly extruded in the villus-like domain. Membrane is labelled with tdTomato. Total duration: 14 hours.

**Supplementary Video 5. Blebbistatin abrogates crypt forces reversibly.**
Organoid monolayer treated with blebbistatin (t=1h 30 min) and allowed to recover after washing out the drug (t=5h). Addition of blebbistatin triggers a rapid reduction of crypt forces. After washout, the crypt recovers its characteristic traction pattern. At t=10h, an epithelial dome is formed. Membrane is labelled with tdTomato. Total duration: 15 hours. Top: 3D traction map of the crypt. Yellow vectors represent components tangential to the substrate and the color map represents the component normal to the substrate. Bottom: lateral view of the organoid monolayer. Yellow vectors represent tractions.

**Supplementary Video 6. Laser ablation between the stem cell compartment and the transit amplifying zone.**
Membrane labelling (left) and velocity field (right) of organoid monolayers before and after ablation. Red line indicates the ablated region. The ablation (t=10s) induces a radial recoil of the monolayer. Membrane is labelled with tdTomato. Total duration: 20s.

**Supplementary Video 7. Laser ablation between the transit amplifying zone and the villus-like domain.** Membrane labelling (left) and velocity field (right) of organoid monolayers before and after ablation (red line). Ablation (t=10s) induces a radial recoil of the monolayer. Membrane is labelled with tdTomato. Total duration: 20s.

**Supplementary Video 8. Cell velocity and traction maps of an intestinal monolayer.**
Cell velocities (left) and 3D tractions (right) of an organoid monolayer. Cell velocity fluctuates in time and space, but shows a general pattern of radial movement away from the crypt. Traction forces are roughly constant in time. Yellow vectors represent components tangential to the substrate and the color map represents the component normal to the substrate. Membrane is labelled with tdTomato. Total duration: 6 hours 30 minutes.

**Supplementary Video 9. Radial laser ablation of a crypt.**
Membrane labelling (left) and velocity field (right) of an organoid monolayer before and after ablation (red line). Radial ablation of the monolayer induces tissue recoil mainly in the direction parallel to the cut. Parallel velocity changes sign at the boundary between the crypt and the villus-like domain. Membrane is labelled with tdTomato. Total time: 20s.
